# Long read-based *de novo* assembly of low complex metagenome samples results in finished genomes and reveals insights into strain diversity and an active phage system

**DOI:** 10.1101/476747

**Authors:** Vincent Somerville, Stefanie Lutz, Michael Schmid, Daniel Frei, Aline Moser, Stefan Irmler, Jürg E. Frey, Christian H. Ahrens

## Abstract

**Background:** Complete and contiguous genome assemblies greatly improve the quality of subsequent systems-wide functional profiling studies and the ability to gain novel biological insights. While a *de novo* genome assembly of an isolated bacterial strain is in most cases straightforward, more informative data about co-existing bacteria as well as synergistic and antagonistic effects can be obtained from a direct analysis of microbial communities. However, the complexity of metagenomic samples represents a major challenge. While third generation sequencing technologies have been suggested to enable finished metagenome-assembled-genomes, to our knowledge, the complete genome assembly of all dominant strains in a microbiome sample has not been shown so far. Natural whey starter cultures (NWCs) are used in the production of cheese and represent low complex microbiomes. Previous studies of Swiss Gruyère and selected Italian hard cheeses, mostly based on amplicon-based metagenomics, concurred that three species generally pre-dominate: *Streptococcus thermophilus*, *Lactobacillus helveticus* and *Lactobacillus delbrueckii*.

**Results:** Two NWCs from Swiss Gruyère producers were subjected to whole metagenome shotgun sequencing using Pacific Biosciences Sequel, Oxford Nanopore Technologies MinION and Illumina MiSeq platforms. We achieved the complete assembly of all dominant bacterial genomes from these low complex NWCs, which was corroborated by a 16S rRNA based amplicon survey. Moreover, two distinct *L. helveticus* strains were successfully co-assembled from the same sample. Besides bacterial genomes, we could also assemble several bacterial plasmids as well as phages and a corresponding prophage. Biologically relevant insights could be uncovered by linking the plasmids and phages to their respective host genomes using DNA methylation motifs on the plasmids and by matching prokaryotic CRISPR spacers with the corresponding protospacers on the phages. These results could only be achieved by employing third generation, long-read sequencing data able to span intragenomic as well as intergenomic repeats.

**Conclusions:** Here, we demonstrate the feasibility of complete *de novo* genome assembly of all dominant strains from low complex NWC’s based on whole metagenomics shotgun sequencing data. This allowed to gain novel biological insights and is a fundamental basis for subsequent systems-wide omic analyses, functional profiling and phenotype to genotype analysis of specific microbial communities.

## Introduction

Metagenomic studies allow the genetic assessment of entire microbial communities. Targeted metagenomic approaches, including the analysis of variable regions of the 16S rRNA, have been widely used to describe the composition of microbial communities [1]. They are particularly useful when a high throughput of samples, deep sequencing of the chosen marker genes and the detection of low abundant taxa is required. However, for a higher resolution assessment of the entire functional potential of microbial communities, whole metagenome shotgun (WMGS) sequencing approaches provide important advantages. They allow researchers to go beyond sequencing and classifying individual genes of species by also covering plasmids, prophages and lytic phages [2], which harbor additional functions and play important roles in shaping microbial communities. Moreover, through the analysis of methylation profiles, one can even link extrachromosomal genetic elements (e.g., plasmids) to their respective host species [3, 4]. Another major objective of WMGS is the resolution of individual strains. This is relevant since specific functions or phenotypic appearances can vary substantially not only between different microbial species, but also among different strains of a species [5]. This functional diversity is derived from genomic variations including larger insertions or deletions resulting in differing gene content, single nucleotide variants (SNV) and varying plasmid content [6]. In order to achieve these key objectives, the assembly of sequencing data needs to be as complete and contiguous as possible. Finished metagenome-assembled genomes (MAGs) harbor more value than near-complete assemblies that still contain gaps [7]. This was illustrated by a recent study, which implicated that long repeat regions of prokaryotic genomes can harbor genes that may confer fitness advantages for the strain [8]. While the major challenge of complete *de novo* genome assembly of individual strains is the resolution of all genomic repeats [8, 9], this situation becomes even more complex for metagenomics: here, the reads do not only have to span intragenomic repeats but also intergenomic repeats, i.e., genomic segments shared by different strains [10]. So far, WMGS studies have mainly relied on short read next-generation sequencing (NGS) technologies, which are not able to span intra- and intergenomic repeats. As a consequence, the assemblies remained highly fragmented [11, 12]. Binning methods, both supervised (reference based) [13] and unsupervised (coverage and nucleotide composition based) [14], have advanced the study of metagenomes to a certain extent [15]. However, it has been suggested that only the use of long range nucleotide information will have the potential to enable complete and contiguous genome assemblies of all dominant species in a microbial community [4, 11]. Recently, such long range nucleotide information including 10X genomics [16], synthetic long-reads [17, 18], Hi-C [11] and very long reads from Pacific Biosciences (PacBio) [19] and Oxford Nanopore Technologies (ONT) [20] have been applied to improve metagenome assemblies. Yet, so far only very few studies have managed to completely assemble genomes without any gaps from complex microbial communities. These included a study of the skin metagenome, in which a single bacterial and one bacteriophage genome could be completely assembled from a complex microbial community using manual curation, while the genomes of a substantial number of co-occurring strains remained in draft status [21]. The proof of concept that it is possible to *de novo* assemble finished MAGs of all dominant taxons in a natural microbial community based on long-read single molecule sequencing data is thus still lacking.

To explore the feasibility of this approach for low complex microbiomes we chose natural whey starter cultures (NWC), which are used in the fermentation step of several types of cheese including Swiss Gruyère. During fermentation, starter cultures from the previous production process are added to the milk, where they metabolize lactose to lactate causing milk acidification. A part of the whey is removed during the cooking process (56-58°C), incubated at 38°C for approximately 20 hours, and subsequently used for the following production batch. As a consequence, whey cultures recurrently encounter considerable environmental changes (e.g., temperature, pH, and redox potential). These harsh culture conditions are known to lead to widespread horizontal gene transfer (HGT) events in the production of fermented foods [22]. Although the NWC microbiomes are of high economic interest, there is limited knowledge on their species and strain composition. This is also true for the presence of plasmids and phages. The latter can have detrimental effects on cheese production if phage-sensitive bacteria are present [23]. Studies performed on NWCs used in the production for Italian hard cheese showed that they contain a low complex lactic acid bacteria (LAB) community. In general, the thermophilic, acid-tolerant, microaerophilic LAB *Streptococcus thermophilus*, *Lactobacillus helveticus*, *Lactobacillus delbrueckii* and *Lactobacillus fermentum* are present [24–27]. The first three species also predominated in a NWC of Swiss Gruyère, as shown by a short read metagenomic approach [28].

The aim of this study was to test the feasibility of *de novo* assembling finished (i.e., complete and contiguous) MAGs from low complex metagenome samples using third generation sequencing data. We hypothesize that we can resolve all dominant strains as well as plasmid and phages, and, thus, gain more meaningful biological insights. Such an approach enables the matching of genotypic and phenotypic characteristics and provide the basis for a subsequent in-depth functional profiling with various omics technologies.

## Results

### *De novo* genome assembly of natural whey culture NWC_1

For NWC_1, we obtained 379,465 PacBio Sequel subreads with an average length of 5,068 bp and a total sequencing output of 1.923 Gb (Additional File 1: Table S1). By using the longest PacBio Sequel reads (147,131 reads >5 kb; 39%), we were able to *de novo* assemble all dominant chromosomes and extrachromosomal elements from this sample. This included two complete, finished circular bacterial genomes, namely *S. thermophilus* NWC_1_1 and *L. delbrueckii* subsp. *lactis* NWC_1_2 (Fig. 1 and Additional File 1: Table S2). The cumulative read output is shown in Additional File 1 (Figure S1). Importantly, we also assembled a matching *L. delbrueckii* subsp. *lactis* plasmid and a matching *Streptococcus* phage (Fig. 1a). Illumina data was only used for polishing steps (see below).

**Fig. 1:**
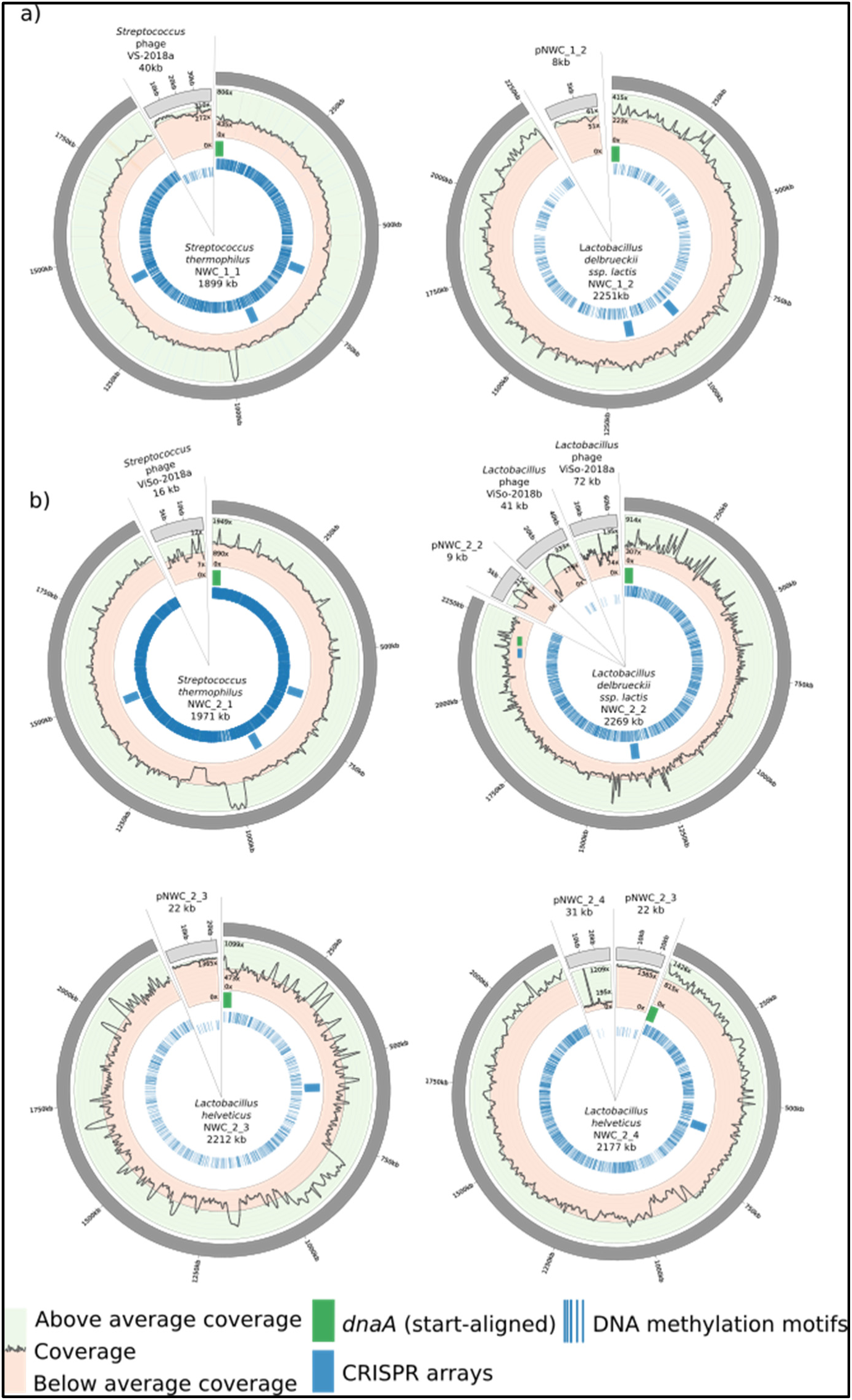
Overview of the genome assemblies of the dominant strains in NWC_1 and NWC_2. a) The Circos plots [30] show the genome assemblies of *S. thermophilus* and *L. delbrueckii* subsp. *lactis, and of a S. thermophilus* phage and the *L. delbrueckii* subsp*. lactis* plasmid from NWC_1 (not drawn to scale), see main text. b) Circos plots are shown for the genome assemblies of *S. thermophilus, L. delbrueckii* subsp. *lactis* and two *L. helveticus* strains from NWC_2, as well as their plasmids and phages (not drawn to scale). The circles illustrate (moving from the outer ring inwards) 1) the genome size, 2) coverage along the genome (green: above average coverage, red: below average coverage), 3) the *dnaA* start point and all CRISPR arrays, 4) all identified DNA methylation motifs that were used to match plasmids to their respective bacterial host.

Maximum likelihood phylogenetic analyses were used to place the newly sequenced strains in the taxonomic context of other finished genomes reported for these species. The average nucleotide identity value (ANIm; calculated from a pair-wise comparison of homologous sequences; m=MUMmer [29]) was used to identify the most closely related strains, plasmids and phages for our *de novo* assembled genomes. The finished *S. thermophilus* NWC_1_1 genome of 1.9 Mbp was characterized by a high sequence coverage (PacBio: 560x, Illumina: 163x) and harbored 2,016 genes including 6 copies of the rRNA operon (Additional File 1: Table S2). It was most similar to *S. thermophilus* APC151 (NZ_CP019935.1; ANIm>99.36; Additional File 1: Figure S3). Similarly, *L. delbrueckii* subsp. *lactis* strain NWC_1_2, also had a high coverage (PacBio: 276x, Illumina: 84x). Its genome was 2.2 Mb in size and contained 2,286 genes including 8 copies of the rRNA operon (Additional File 1: Table S2). It was most similar to *L. delbrueckii* subsp*. lactis* DSM 20072 (ANIm>99.22; Additional File 1: Figure S4). Moreover, the circular plasmid pNWC_1_2 (8kb, 9 genes) was most similar to plasmid pLL1212 (ANIm>96.01), which was originally isolated from *L. delbrueckii* subsp. *lactis* (Genbank AF109691). The assembly of the complete, linear *Streptococcus* phage VS-2018a genome (40kb, 55 genes) was most similar to *Streptococcus* phage TP-778L (ANIm>91.47).

Importantly, overall, 99.3 % of the quality-filtered Illumina reads mapped back to these assemblies. This indicated that we managed to assemble the most dominant (relying on >1% of Illumina reads as arbitrary cut-off), and thus, presumably most relevant species of this microbial community.

### De novo genome assembly of natural whey culture NWC_2

Relying on PacBio Sequel data alone, we were not able to completely assemble all dominant genomes from NWC_2, indicating that its complexity, i.e., the number of dominant species and strains, was higher than that of NWC_1. Binning of the PacBio pre-assembled reads did not completely disentangle the genomes. Neither for NWC_1 (Additional File 1: Figure S6; carried out retrospectively for comparison) nor for NWC_2 (Additional File 1: Figure S7) could we distinguish the dominant prokaryotic genomes present based on their coverage (supervised, abundance based binning), nor their GC content or tetranucleotide frequency (unsupervised, compositional binning). While some binning methods worked to a certain degree for NWC_1 (Additional File 1: Figure S6d) and for NWC_2 (Additional File 1: Figure S7c), no method was able to bin all pre-assembled reads into the appropriate species bin and thereby avoid “contamination” (i.e., reads from other genomes). Furthermore, we observed that two contigs (phage NWC_2_1, pNWC_2_2) were not covered by any pre-assembled PacBio read (see Additional File 1: Figure S7, legend). This is most likely due to the fact that for the pre-assembly only the longest reads are considered, whereby shorter extrachromosomal contigs (e.g., phages and plasmids) are statistically less often considered.

We therefore also generated ONT data for this sample, aiming to use the longest reads for the assembly. We obtained 407,027 ONT reads with a total sequencing output of 1.385 Gb (Additional File 1: Table S1 and Figure S2). A cumulative read output analysis of both PacBio and ONT data indicated that -in theory-we should now be able to span the longest repeats with the ONT data (Additional File 1: Figure S2). By using long ONT reads from NWC_2 (>20kb; longest mappable read: 118,642 bp), we were finally able to *de novo* assemble finished MAGs of all dominant species and strains. Remarkably, this included two distantly-related strains of the same species (*L. helveticus*). Overall, we assembled four finished bacterial genomes including *S. thermophilus* strain NWC_2_1 and *L. delbrueckii* subsp. *lactis* strain NWC_2_2, two *L. helveticus* strains NWC_2_3 and NWC_2_4, and finished versions of three plasmids and three phage genomes (Fig. 1b, Additional File 1: Table S2). Illumina data was used for polishing steps (see below).

High coverage was achieved for the complete *S. thermophilus* NWC_2_1 genome (ONT: 160, PacBio: 833, Illumina: 69; Additional File 1: Table S2), which was most similar to *S. thermophilus* APC151 (NZ_CP019935.1; ANIm>99.35; Additional File 1: Figure S3). The genome of 1.9 Mb harbored 2,108 genes including 6 copies of the rRNA operon. For this genome, we could also identify a corresponding *Streptococcus* phage ViSo-2018a (see below; 15kb, 15 genes), which was most similar to *Streptococcus* phage P9854 (KY705287.1; ANIm>98.74). Furthermore, the *L. delbrueckii* subsp. *lactis* NWC_2_2 genome (ONT: 63x, PacBio: 273x, Illumina: 54x) of 2.2 Mb which encoded 2,331 genes including 8 copies of the rRNA operon (Additional File 1: Table S2) was most similar to *L. delbrueckii subsp.* lactis DSM 20072 (ANIm>99.16; Additional File 1: Figure S4). For this strain, we were able to identify one matching plasmid pNWC_2_2 (9kb, 8 genes) with high coverage (ONT: 160x, PacBio: 833x and Illumina: 69x; Additional File 1: Table S2), that was most closely related to plasmid pLL1212 (ANIm>96.02). For the phage genomes, we could identify that *Lactobacillus* phage ViSo-2018b (41kb, 86 genes) was most closely related to *Lactobacillus* phage phiJB (ANIm>87.25) and *Lactobacillus* phage ViSo-2018a (72kb, 85 genes) to *Lactobacillus* phage Ldl1 (ANIm>97.51). Importantly, we were able to disentangle the two *L. helveticus* NWC_2_3 and NWC_2_4 strains. They harbored 2,385 and 2,318 genes respectively, with 5 RNA operon copies each (Additional File 1: Table S2). They were most similar to *L. helveticus* FAM8627 (ANIm=99.63) and FAM8105 (ANIm=99.57; Additional File 1: Figure S5). Further, we assembled two circular plasmids. Plasmid pNWC_2_3 (22kb, 21 genes) was most similar to pL11989-1 (ANIm>94.84) and pNWC_2_4 (31kb, 29 genes) most similar to plasmid H10 (ANim>94.58).

The extensive polishing of the assemblies with all available sequencing data was crucial for the generation of finished high quality genomes, especially for the more complex NWC_2 sample (Additional File 1: Figures S8 amd S9, Additional File 2). Using an iterative polishing approach, we were able to continuously reduce misassemblies (Additional File 1: Figure S8a) by removing mismatches and indels (Additional File 1: Figure S8b) and thereby increasing the covered fraction compared to the finished genome sequence (Additional File 1: Figure S8d). In addition, the pseudogene count can serve as a quality measure for third generation sequencing based genome assemblies [31]. Overall, we observed a decrease of the total number of pseudogenes over the course of the polishing steps. The pseudogene counts for the final polished genome sequences were comparable to those reported for other strains of the respective species (Additional File 1: Figure S9c, Table S3, Additional File 2). Importantly, 99.0 % of the quality-filtered Illumina reads could be mapped back to the MAGs. This suggested that we could also assemble the genomes of all dominant species and strains of this microbial community.

### Advantages of complete PacBio/ONT assemblies over fragmented Illumina assemblies

To illustrate the advantages of our long read-based finished MAGs, we compared the PacBio/ONT bacterial assemblies versus the respective Illumina-only based metagenome assemblies (Fig. 2). For NWC_1 and NWC_2, we obtained 2,132,096 and 1,410,764 Illumina reads (300 bp PE), respectively, of which the large majority (94% and 93%, respectively) was of high quality and paired (see Additional File 1: Table S1). An assembly of the Illumina data using metaSPAdes [32] resulted in highly fragmented assemblies for both metagenome samples (Fig. 2a,b; track 2). The Illumina assemblies were characterized by a much lower contiguity, i.e., larger number of contigs (NWC_1: 2,452 contigs, NWC_2: 4,524 contigs) and covered only ~88% and ~66% of the NWC_1 and NWC_2 genome sequences, respectively (Fig. 2a,b: track 3).

**Fig. 2:**
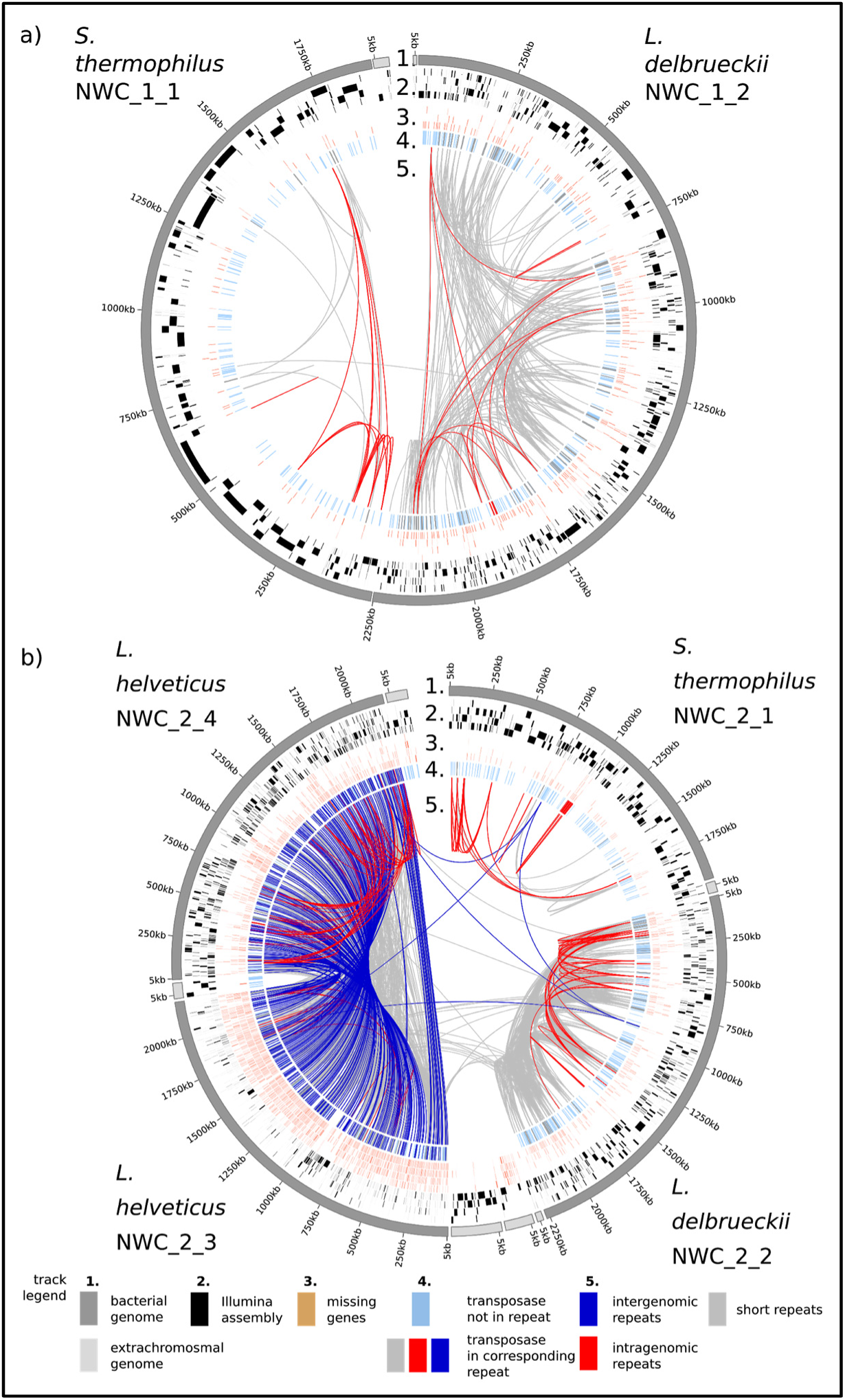
Comparison of complete PacBio/ONT and fragmented Illumina assemblies for a) NWC_1 and b) NWC_2. Description of tracks from outer towards inner tracks: 1) All completely assembled contigs (plasmids and phages in light gray) as reference. 2) The Illumina assembled contigs mapped to the reference. 3) Genes that are missing in the respective Illumina assemblies. 4) Transposases that are either located in repeat regions (dark blue) or not (light blue). 5) Intragenomic (red) and intergenomic repeats larger than 3 kb and 95 % identity (blue) and short repeats (> 1.5 kb, > 3kb) and 90 % identity (gray).

A large percentage of the assembly breaks can be explained by repeat regions occurring within (intragenomic) or between (intergenomic) the genomes (Fig. 2a,b; track 5.). These intra- and intergenomic repeats consisted mainly of multicopy genes (e.g., transposases) or of conserved regions (e.g., rRNAs) called synteny blocks [10] (Fig. 2a,b; track 4.). Lactobacillus in general [33], and our assemblies in particular (Additional File 1: Table S3), contain large numbers of transposases which account for a substantial part of these intra- and intergenomic repeats (95% and 81% for NWC_1 and NWC_2, respectively) (Fig. 2 track 5). Overall, the Illumina assemblies resulted in lower quality genome annotations for the bacterial strains of NWC_1 and NWC_2, affecting roughly 11 % (397 of 3,644) and 37 % (2,785 of 7,451) of the annotated genes, respectively (Fig. 2 track 3). The intergenomic repeats become more problematic when several strains of a species are present in the metagenome sample as we can observe in NWC_2 (Fig. 2 track 5.).

### 16S rRNA taxonomic profiling supports the long read-based assembly results

We independently assessed the community composition of the two NWCs using a 16S rRNA amplicon-based approach and compared it to metagenomic taxon profiling of Illumina and PacBio data (full details can be found in Additional File 1: Tables S5 and S6, Figures S10 and S11). Oligotyping of the 16S rRNA amplicon data resulted in the delineation of 3 dominant oligotypes overall, which could be identified on the species level (Fig. 3), and 6 very low abundant oligotypes, which could be identified either on the species or genus level (Additional File 1: Table S5). *S. thermophilus* was the dominant species in both samples with a relative abundance of 65.4% in NWC_1 and 45.4% in NWC_2. *L. delbrueckii* was the second most abundant species with a relative abundance of 34.1% in NWC_1 and 24.5% in NWC_2. *L. helveticus* made up 0.1% of the community in NWC_1 and 25.6% in NWC_2. A rarefaction analysis of these data resulted in plateauing curves (Additional File 1: Figure S10), which indicated that the large majority of species was found. Similar results were obtained from the compositional estimations based on an analysis of the Illumina reads using Metaphlan2 [34] and of the PacBio reads using MetaMaps [35]. Compared to the other two analyses methods, the MetaMaps analysis of PacBio reads resulted in a somewhat elevated percentage of reads that could not be assigned to taxa and to a higher/lower abundance of *L. helveticus*/*L. delbrueckii* in NWC_2 (Fig. 3, Additional File 1: Table S6).

**Fig. 3:**
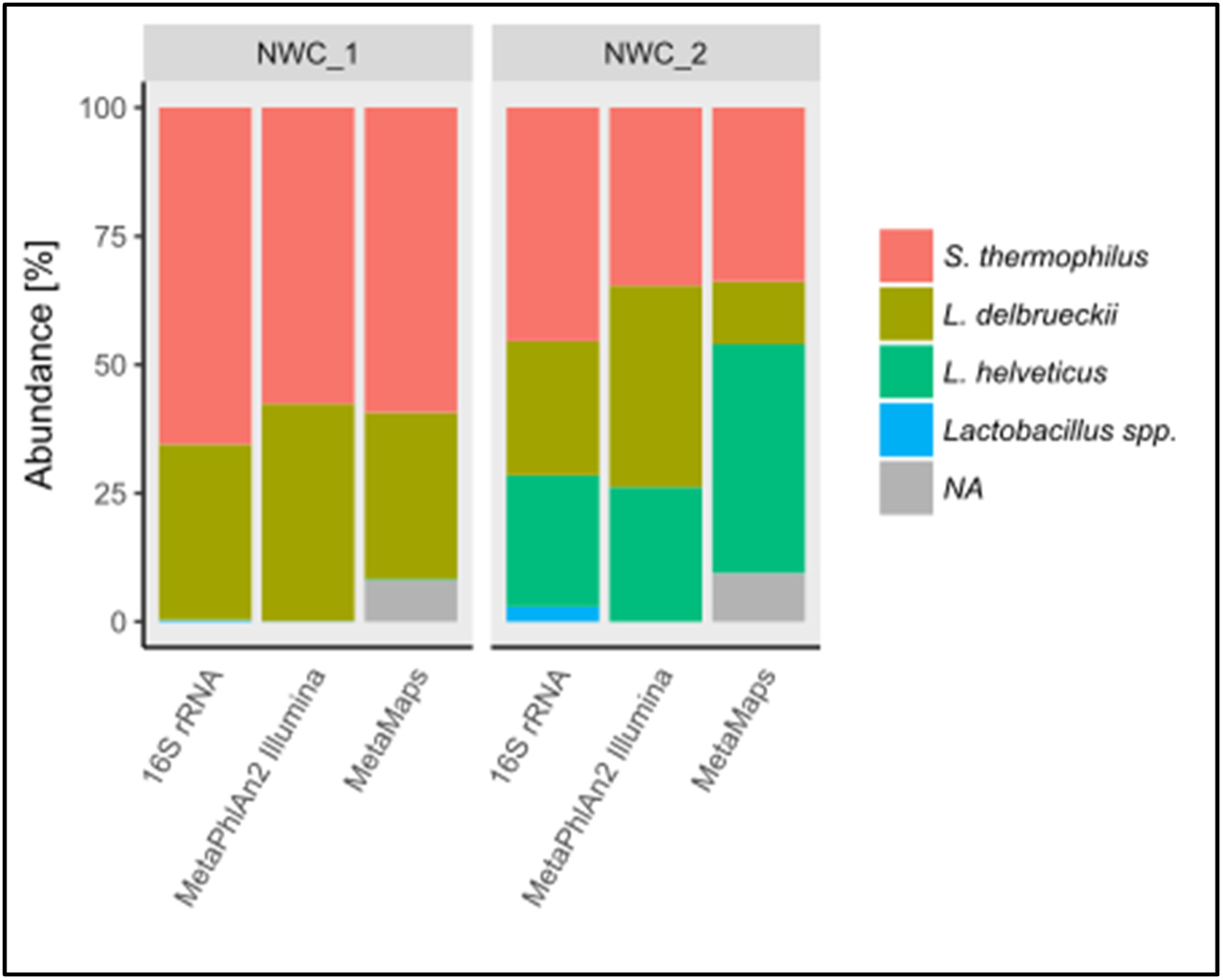
Taxonomic profiling of NWC_1 and NWC_2. The relative abundances of predominant species in NWCs (see legend) are based on the 16S rRNA (v4) amplicon data, a Metaphlan2 [34] analysis of the Illumina data, and a MetaMaps [35] analysis of the PacBio data for NWC_1 and NWC_2, respectively. NA=not assigned.

### Resolution of the two assembled *L. helveticus* strains in NWC_2

The co-assembly of two unique *L. helveticus* strains in NWC_2 was achieved by extensive polishing of a scaffolded assembly combined with a more detailed coverage analysis. The initial *de novo* assembly based on ONT reads resulted in 12 scaffolded *L. helveticus* contigs. From the assembly graph, we could infer that two circular *L. helveticus* strain genomes were present, which were clearly distinct over the majority of their genomes (3.833 Mb of 4.063 Mb, 94%; Fig. 4a). However, four regions remained, which could not be completely spanned with the current sequencing output. Yet, based on the coverage of the individual contigs we could separate the contigs into a low (~30x) and high (~60x) coverage strain (Fig. 4b), while the “shared” contigs roughly exhibited coverage of ~90x (i.e., similar to the summed coverage). Even genome coverage was observed at the locations where the contigs were merged (Fig. 4e and f). Overall, this indicates the correct assembly of the two genomes. The *L. helveticus* strain identity and abundance was also analyzed by high-throughput *slp*H amplicon sequence typing [36] (Additional File 1: Figure S11). The two dominant sequencing types ST13 (74%) and ST38 (19%) corresponded in both abundance (NWC_2_4: 69.9%, NWC_2_3: 30.1%; Fig. 4c) as well as sequence identity to the *slp*H sequences extracted from the assembled *L. helveticus* strains NWC_2_3 and NWC_2_4, and were in par with the abundance values estimated by MetaMaps (Fig. 4d). Finally, when aligning the genomes of the two putative *L. helveticus* strains against each other, major genomic rearrangements were revealed (Fig. 4g). In addition, the two genomes shared 1,258 genes (core genes) and contained 555 (NWC_2_3) and 525 (NWC_2_4) unique genes. Among the unique genes, the large number of transposases (category L, “Replication, recombination and repair”) was striking. In addition, the unique genes of *L. helveticus* NWC_2_3 were enriched for “Nucleotide transport and metabolism” and those of *L. helveticus* NWC_2_4 for “Defense mechanisms” (Additional File 1: Table S7). Overall, this is well in line with their separate placement on a phylogenetic tree built from all finished *L. helveticus* genomes (see Additional File 1: Figure S5).

**Fig. 4:**
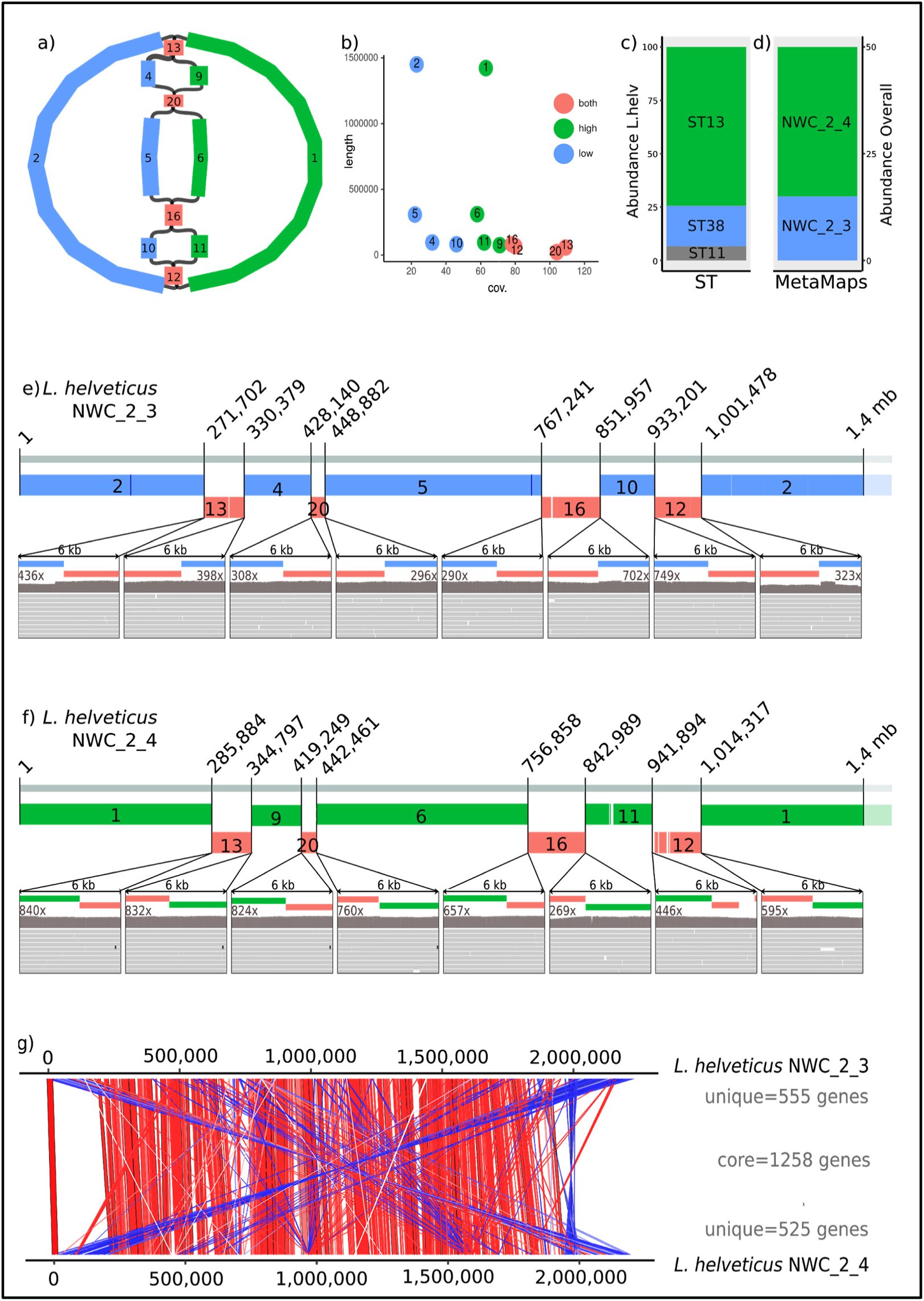
Resolution of two distantly related *L. helveticus* strains in NWC_2. a) Assembly graph from Bandage [37] colored according to high (green) or low (blue) coverage contigs as well as genomic regions that occur in both strains (red) before genome polishing. The numbers correspond to the respective contigs visualized in b). b) Coverage plot of the individual contigs. c) Abundance of *L. helveticus* sequence types based on *slpH* sequence typing. d) *L. helveticus* abundance based on PacBio coverage. e) PacBio reads spanning the initial contig gaps after polishing of *L. helveticus* NWC_2_3 and f) *L. helveticus* NWC_2_4. g) Synteny plot of *L. helveticus* NWC_2_3 and NWC_2_4 with the number of core and unique genes. Regions of similarity are indicated by red (same orientation) and blue (opposite orientation) bars.

### Matching plasmids to host strains

As plasmids do not contain methyltransferases, their DNA methylation is determined by the host [38]. Therefore, DNA methylation motif detection allowed us to match plasmids and host genomes. For NWC_1, we could detect DNA methylation motifs in both bacterial chromosomes (Additional File 1: Figure S12). However, due to the low read coverage and likely also its small size, we were not able to identify a DNA methylation motif on plasmid pNWC_1_2 (Fig. 1, Additional File 1: Figure S12). Nevertheless, this plasmid was most closely related to the previously sequenced *L. delbrueckii* subsp*. lactis* plasmid pLL1212 (Genbank AF109691; ANIm>96.01). For NWC_2, we were able to assemble three plasmids. One plasmid (pNWC_2_2) was highly similar to plasmid pNWC_1_2/pLL1212; as already observed for NWC_1, we could not detect a methylation motif either (Fig. 5). For the other two plasmids, we could identify DNA methylation motifs that matched motifs uniquely occuring in *L. helveticus* (Fig. 5). Based on the coverage of the plasmids, we suggest that plasmid pNWC_2_4 only occurs in *L. helveticus* strain NWC_2_4, while the second plasmid pNWC_2_3 likely occurs in both *L. helveticus* NWC_2_3 and NWC_2_4 strains.

**Fig. 5:**
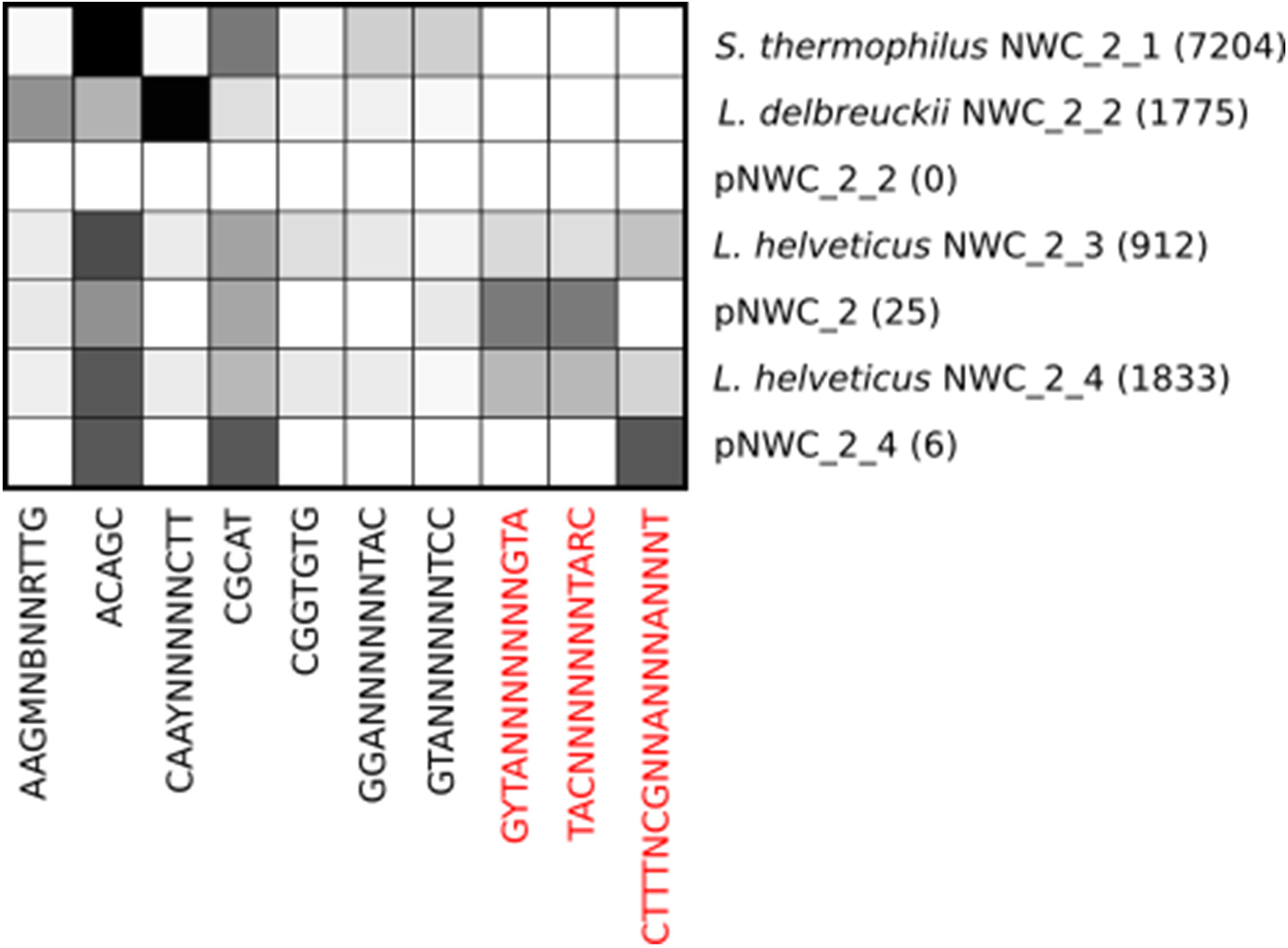
DNA methylation motif analysis. The sequence and abundance of DNA methylation motifs were determined in all *de novo* assembled genomes of NWC_2 with the base modification module of the SMRTlink (v.5.1.0) toolkit and visualized. The heatmap illustrates the relative abundances of the motifs per assembly (increasing relative abundance from white to black). The numbers in the brackets represent the number of DNA methylation motifs detected in a given assembly. Motives specific to the *L. helveticus* strains and plasmids are highlighted in red.

### Matching CRISPR arrays and targets

Matching CRISPR arrays present in bacterial genomes and protospacer sequences in phage genomes can help to explain the susceptibility of the strains to the phages present in a metagenome sample [39]. We were able to identify several CRISPR arrays in all bacterial genomes of NWC_1 and NWC_2 (Fig. 1, Additional File 1: Table S8). For six CRISPR spacers in two CRISPR arrays of *S. thermophilus* NWC_1_1, we found closely matching (less than three mismatches among the roughly 30 bp spacer sequence) protospacer sequences in the assembled phage genome (Fig. 6). This suggests a previous encounter of this phage with *S. thermophilus* strain NWC_1_1, indicating an acquired resistance of the bacterium against this phage. Further, we were able to identify five different Cas protein-coding genes in proximity of the CRISPR arrays of *S. thermophilus* NWC_1_1 (Fig. 6). Overall, this indicates that the CRISPR arrays are still active.

**Fig. 6:**
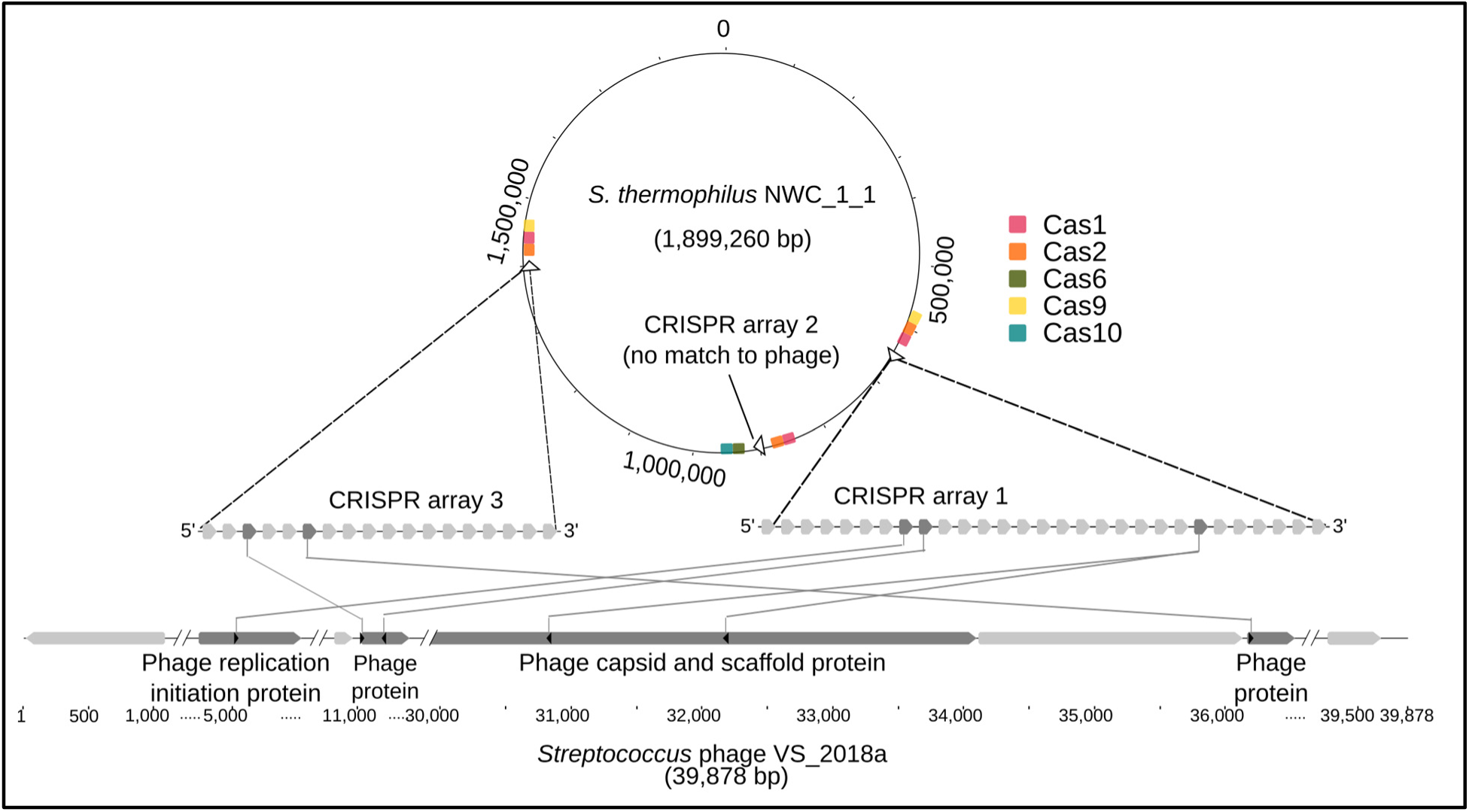
CRISPR spacers in *S. thermophilus* strain NWC_1_1 and the *S. thermophilus* phage genome. Three CRISPR arrays (open arrows) and their flanking Cas genes are shown in the genome of strain *S. thermophilus* NWC_1_1 (top). CRISPR arrays 1 and 3 have matching spacers with the phage, as shown in the zoomed regions of the ~40 kb phage genome along with the annotation of selected phage protein-coding genes (bottom).

Similarly, matches of CRISPR arrays and protospacers were found for strain *S. thermophilus* NWC_2_1 and *Streptococcus* phage ViSo-2018a (four matches) and for *L. delbrueckii* subsp. *lactis* NCW_2_2 and *Lactobacillus* phage ViSo-2018a (four matches). However, for strain *L. delbrueckii* subsp. *lactis* NWC_2_2 and the *Lactobacillus* phage ViSo-2018b only a single match with six mismatches to the spacer sequence was found. The relatively poor match of a CRISPR spacer and the phage protospacer could potentially indicate a diminished protection against a corresponding phage. This might result in a partial susceptibility of *L. delbrueckii* subsp. *lactis* NWC_2_2 to *Lactobacillus* phage ViSo-2018a and explain the high coverage of the *Lactobacillus* phage ViSo-2018a. Similarly, the *S. thermophilus* prophage has only a single low quality (five mismatches) match with the CRISPR spacer sequence in the *S. thermophilus* NWC_2_1 genome (Additional File 1: Table S8).

### Genome comparison of the two *S. thermophilus* strains reveals the presence of an active phage

The genomes of the two *S. thermophilus* strains from NWC_1 and NWC_2 shared a very high amount of sequence identity (ANIm>99.7%). Overall, 88 variants (71 SNPs, 5 insertions and 12 deletions) could be detected between the two annotated genomes. Notably, we identified two larger insertions in the genome of *S. thermophilus* NWC_2_1 compared to NWC_1_1. The first insertion represented a triplet tandem repeat of the extracellular polysaccharides (EPS) type VII operon, i.e., 2 additional copies of the operon compared to strain NWC_1_1 (Additional File 1: Figure S13). The second insertion could be linked to an inserted prophage (41kb, 55 annotated genes, see Fig. 7). We observed reads which mapped both to the bacterial genome and extending into the prophage genome and *vice versa* (Fig. 7b), providing proof of the integration into the bacterial host genome. This variant was supported by approximately 22% of the reads at the prophage start position. However, the majority of reads (71%) mapped to the bacterial genome without the sequence of the putative prophage (Fig. 7c). Further, we also encountered a substantial amount of reads (n=47, 7%) that spanned over the end of the prophage genome and back into the reverse opposite end of the prophage (Fig. 7d). This suggested that a certain fraction of the phage genome is circular and was therefore also occurring in a non-inserted (i.e., lytic) state. Further, the *S. thermophilus* genome did not harbour any CRISPR array spacers that matched the prophage. We also observed that the prophage inserted just upstream of a tRNA-Arg. Overall, we assume this to be an example of an active phage system.

**Figure 7:**
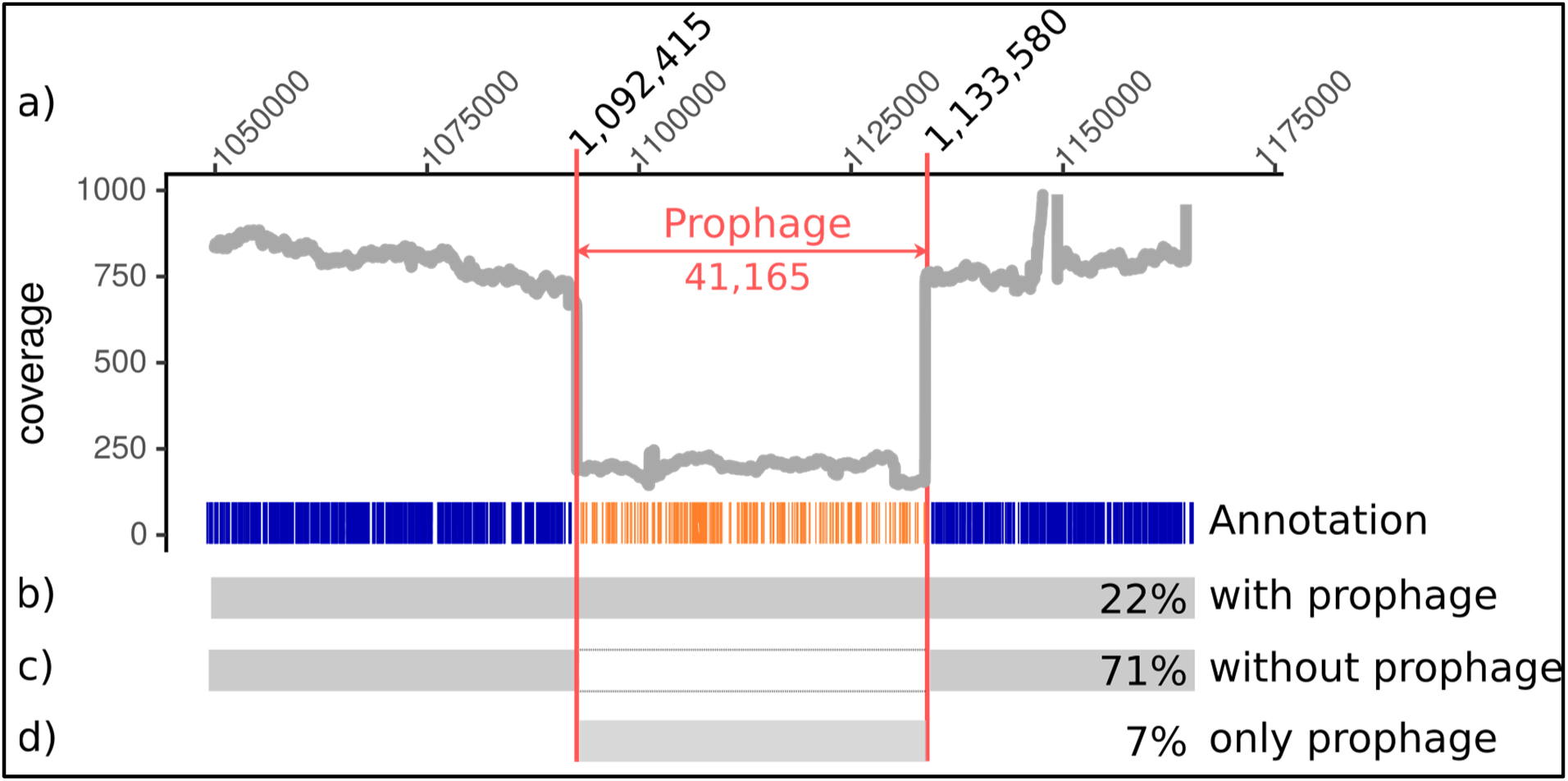
Genome coverage of *S. thermophilus* NWC_2_1 around the prophage insertion site. a) Genome coverage of *S. thermophilus* NWC_2_1 in the proximity of the prophage. Below the coverage plot, we highlight the prophage genome and its annotation as well as the percentage of reads that supported a respective variant. b) The bacterial genome variant with the inserted prophage. c) The dominant bacterial genome variant without the prophage. d) The phage variant (here shown as linearized genome).

## Discussion

In this study, we demonstrated the feasibility of complete *de novo* genome assembly of all dominant species, directly from low complex metagenomes using third generation long-read sequencing. This included the resolution of two distinct strains of *L. helveticus* in one sample and the recovery of several plasmids and phage genomes. Furthermore, by matching methylation patterns as well as CRISPR arrays and protospacer elements, we could link several of the observed plasmids and phages with their respective bacterial hosts and uncover evidence for previous encounters between bacterial strains and phages.

The read length of third generation sequencing technologies (i.e., PacBio and ONT) was instrumental to achieve finished assemblies of the dominant MAGs. So far, a number of studies have reported the recovery of genomes from highly complex metagenomes [21, 40], which were, however, predominantly based on the assembly of short reads, and thus, did not represent finished genomes [41, 42]. With the “Illumina only” assemblies, we could illustrate that they missed a significant percentage of genome regions which could be covered by finished MAGs based on long reads (Fig. 2). Binning, a common approach to assign short metagenomic reads from complex samples to their respective genomes before assembly, aims to take advantage of differences in coverage [43], tetranucleotide frequency [44] or GC content. However, complete binning of pre-assembled PacBio reads could not be achieved in our study, despite the low number of species, long read data and divergent GC content between the genomes. Several reads were not clearly separated (Additional File 1: Figure S6 and S7), which could partially be attributed to the low average read length of the PacBio Sequel reads. Further method development on the sample processing aspects and sequencing technology is expected to provide even longer fragments with lower error rates. For our low complex samples, the higher error rates of third generation sequencing technologies could be removed as a sufficiently high sequencing coverage was achieved. Longer reads should eventually be able to overcome the need for binning approaches even in more complex microbial communities.

To our knowledge, there are currently no well-established long-read metagenome assemblers available or they are still in an experimental state (e.g., Flye-meta). Here, we used the Flye *de novo* assembly algorithm [45], which was initially developed for individual repeat rich genomes, yet, achieved the best assemblies of our metagenomic samples (data not shown). Further, it was crucial to extensively polish genome assemblies in order to achieve a sufficiently high assembly quality [31] (Additional File 1: Figure S8, Additional File 2). We found that very long reads (ONT) were necessary to resolve long-range mis-assemblies. However, the lower quality of ONT reads required polishing with PacBio and in particular Illumina data. Moreover, great care needs to be taken when contigs are polished individually, since this can lead to the erroneous removal of true, natural sequence diversity due to cross mapping of reads in repeat regions (e.g., repeated sequences such as 16S rRNA operons, insertion sequences/transposases). Furthermore, we still observed a high number of pseudogenes in the finished MAGs. This, however, is characteristic for *Lactobacillales*, which live in a nutrient-rich environment such as milk and therefore frequently experience gene loss and gradual genome decay [28, 46]. Overall, further improvements of the sequencing technologies (ONT/PacBio), the application of long-range information technologies (e.g., 10x genomics, Hi-C, synthetic long reads) combined with the development of new algorithms could greatly simplify the currently extensive assembly and polishing workflow.

The identification of taxa in an assembled metagenome and the estimation of their abundance is often the first step of a microbial community analysis. Many taxon profilers exist for Illumina shotgun metagenomics data [47]. However, due to the intrinsic differences in quality and read length, these methods are not transferable to long reads. Only a few very recently developed taxon profilers can cope with long reads, such as MEGAN-LR [48] and MetaMaps [35]. We individually assessed taxa abundance based on WMGS PacBio (MetaMaps) and Illumina (Metaphlan2) data, as well as a targeted amplicon approach using the v4 region of the 16S rRNA. The abundance values of the strains based on the PacBio based MetaMaps approach were not entirely in par with the findings derived from the 16S rRNA amplicon and Illumina based Metaphlan2 approach (Fig. 3, Additional File 1: Table S6). Almost 10% of the PacBio reads in each sample could not be assigned to taxa (Additional File 1: Table S6). This could be due to sequencing errors in low quality sequences, and thus, no matches in the reference database. To a certain extent, the differences could also be caused by abundance biases introduced in the PacBio library preparation process, either by unequal shearing of genomic DNA by the Megaruptor device, or during the enrichment for long fragments. The original abundance ratios are thus likely best reflected in the Illumina data, in particular since more than 99% of the reads could be mapped to the finished MAGs, plasmids and phage genomes. All dominant strains identified her accounted for at least 1% of the Illumina reads.

Within undefined cheese starter culture communities there are usually multiple strains per species with only a few being dominant [49]. Our long read-based approach could identify all dominant members of the community and the targeted survey based on 16S rRNA amplicon data resulted in the detection of only a few, additional very low abundant taxa, which are presumably of minor importance in our samples. Most importantly, our approach enhanced the taxonomic resolution down to the strain level for the most dominant strains, which represents a significant advantage over other approaches. Interestingly, the strains identified in the NWCs from two different cheese producers included examples of almost identical genomes (for the *S. thermophilus* strains; see below), moderately different genomes for the *L. delbrueckii* strains (1608 core genes, 110 and 152 strain-specific genes) up to quite distinct *L. helveticus* strains co-occurring in the same sample (1300 core genes, 555 and 525 strain-specific genes). This clearly illustrates the value of assembling complete genomes as the strains might harbor substantial functional differences beyond the reach of amplicon based methods. Furthermore, our results show that the complexity of our NWC metagenome samples was even lower than implied by previous studies [50]. The absence of *L. helveticus* in NWC_1 was particularly striking, since this species is thought to play an essential role in the production of Swiss Gruyère [49, 51]. The presence of *L. helveticus* strains results in the reduction of the cheese bitterness (due to their proteolytic activity) [52], as well as in a faster ripening and enhanced flavor development, which are desirable effects in the production of cheese [53, 54]. Yet, in certain production steps their activity can also lead to undesirable effects including the formation of splits and cracks and reduced elasticity due to an excessive proteolysis and carbon dioxide production [55]. Since *L. helveticus* is thought to be more heat sensitive compared to the other predominant NWC species, this might in part explain the reduced diversity in NWC_1 at the time of sampling. For biotechnological applications, it is necessary to differentiate and characterize the different strains. Strain typing has been of major interest in many fields of microbiome research [56]. Sophisticated tools such as PanPhlAn [57] or mOTU [58] have been developed to circumvent an assembly and reveal strain diversity from raw Illumina data. However, such approaches are limited since they rely on reference databases. Here, we show an alternative approach by using long read information. With increasing community complexity, the strain resolution becomes more tedious, as was the case for NWC_2. Yet, we were able to assemble two finished genomes of two strains of the same species (i.e., *L. helveticus*, Fig. 4), and thus, gain the complete genomic information of the strains present.

In contrast to *L. helveticus*, *S. thermophilus* and *L. delbrueckii* subsp. *lactis* were present in both NWC metagenome samples and are known to exist in tight association [59]. *S. thermophilus* actively supports *L. delbrueckii* subsp. *lactis* growth by producing acid and dissolving oxygen to CO_2_, thereby creating the optimal anaerobic conditions necessary for *L. delbrueckii* subsp. *lactis* to thrive. In return, *L. delbrueckii* subsp. *lactis* stimulates *S. thermophilus* growth by the release of amino acids through proteolytic enzymatic activity [60]. The two *S. thermophilu*s strains assembled from NWC_1 and NWC_2 shared a high amount of sequence identity, yet, their comparison revealed intriguing genomic differences including the insertion of two additional repeats of the eps operon in strain NWC_2_1 compared to strain NWC_1_1 (Additional File 1: Figure S13). The synthesis of extracellular polysaccharides is widespread in many *S. thermophilus* strains [61]. EPS production can impart a positive effect on the functional properties of cheese (i.e., texture, viscosity) [62, 63]. Furthermore, capsular EPS are thought to protect bacteria against detrimental environmental conditions including phage attacks [62]. Yet, so far this has not been shown for LAB, and thus, cheese producers cannot solely rely on the EPS production of *S. thermophilus* to protect starter cultures against phage infections. EPS in *S. thermophilus* strains are known to vary considerably in their repeating structures [62], which was also the case for our assembled strains. These genes would represent interesting candidates for subsequent genotype to phenotype analyses, i.e., to explore whether strain-specific differences in EPS production could affect their protection potential against phages. This could have practical applications, as phages can cause failures in the fermentation process and result in severe economic losses to the cheese industry [64].

On the other hand, phages can likely act as vectors for horizontal gene transfer, which is a common phenomenon in the dairy production [22]. Here we could uncover evidence for such an active phage system by assembling the bacterial host genome, as well as the inserted prophage and lytic phage. Moreover, past encounters of phages and bacteria could be revealed by the matching of protospacers in the bacteriophage and clustered regularly interspaced short palindromic repeats (CRISPR) in the bacterial genome, which represent an acquired immunity [65] [66]. Here we were able to assemble four complete phage genomes with matching CRISPR arrays. Interestingly, the assembled genomes in NWC_2 did not show good CRISPR matches with the most abundant phage (*Lactobacillus* phage ViSo-2018a) and the prophage inserted in *S. thermophilus* NWC_2_1. This might indicate that the occurring CRISPR spacers are inefficient in providing protection against the phages.

Finally, another crucial advantage of finished MAGs is the possibility to associate plasmids with their most likely bacterial host. Currently, only PacBio and ONT are able to detect DNA methylation motifs by sequencing, which allowed us to match four circular plasmids with their respective bacterial host species. The complete genome information encompassing the genes on chromosome and plasmid(s) provides the basis for a systems-wide functional profiling and the potential discovery of important genes coding for antibiotic resistance [67], virulence factors [68] or specific traits that are beneficial for cheese production [69], which was, however, beyond the scope of this study.

## Conclusions

Relying on long reads from third generation sequencing technologies, we demonstrate the feasibility of *de novo* assembling finished MAGs for the dominant strains from cheese starter cultures, which represent low complex metagenomes. Of particular value were the insights gained from the assembly of co-occurring prophages, phages and plasmids, which uncovered evidence of previous bacteriophage encounters and contributed to the comprehensive assessment of the overall functional potential of these microbial communities.

## Supporting information

## Methods

### NWCs and genomic DNA isolation

NWCs were collected at two Swiss Gruyère cheese PDO factories at the time of cheese production (four 50 mL aliquots per sample) and transferred to the lab on ice. Genomic DNA (gDNA) was immediately isolated by mixing each sample aliquot with 0.25 mL of 10% (w/v) sodium dodecylsulfate and centrifugation (30 min at 20 °C, 4,000 g). The supernatants were removed leaving a volume of 5 mL to resuspend the pellet. After pooling suspensions of the same NWC sample, aliquots of 1 mL were centrifuged at 20°C for 5 min at 10,000 g, supernatants were discarded and gDNA was extracted from the pellets as previously described (Moser et al., Int. Dairy J. 2017).

### PacBio Sequel library preparation, WMGS sequencing and read filtering

The SMRTbell was produced using PacBio’s DNA Template Prep Kit 1.0 as follows: input gDNA concentration was measured with a dsDNA Broad Range assay on a Qubit Fluorometer (Life Technologies); 10μg of gDNA were sheared mechanically with a Megaruptor Device (Diagenode, Liege, Belgium) to an average fragment size distribution of 15-20 kb, which was assessed on a Bioanalyzer 2100 12Kb DNA Chip assay (Agilent). Five μg of sheared gDNA were DNA damage repaired and end-repaired using polishing enzymes. A blunt end ligation reaction followed by exonuclease treatment was performed to create the SMRTbell template. A Blue Pippin device (Sage Science) was used to size select the SMRTbell template and enrich for fragments > 10 Kbp. The sized selected library was quality inspected and quantified on an Agilent Bioanalyzer 12Kb DNA Chip and on a Qubit Fluorometer, respectively. A ready to sequence SMRT bell-Polymerase Complex was created using PacBio’s Sequel binding kit 2.0 according to the manufacturer’s instructions. Each sample was sequenced on 1 Sequel™ SMRT^®^ Cell 1M v2, taking a 10 hour movie using the Sequel Sequencing Kit 2.1. The sequencing data quality was checked via PacBio’s SMRT Link (v5.0.1) software, using the “run QC module”. As the sequencing data from the Sequel platform (v.2.1) does not provide a read quality score nor a per base quality score, metrics that otherwise can guide the selection of an optimal subset for a *de novo* genome assembly, read selection was based on read length. To allow assembly of the dominant genome variant(s) of the present species, we filtered the NWC_1 data for the longest reads (> 5 kb, n=147,131).

### Oxford Nanopore library preparation, WMGS sequencing and read filtering

For NWC_2, an additional ONT library was prepared using a 1D2 Sequencing Kit (SQK-LSK308) and sequenced on a FLO-MIN107 (R9.5) flow cell. In order to assemble the dominant genome variant(s) of the present taxa, base called reads were filtered for reads > 20 kb (n=32,829) using Filtlong v.0.2.0. In addition, we discarded the 10% of lowest quality reads based on their Phred quality scores.

### Illumina MiSeq library preparation, WMGS sequencing and read filtering

Two 2 × 300 bp paired end libraries were prepared per sample using the Nextera XT DNA kit and sequenced on a MiSeq. The reads were paired with trimmomatic (v0.36); only paired reads were used for the final mapping (parameters: “LEADING:3 TRAILING:3 SLIDINGWINDOW:4:15 MINLEN:36”). A subset of the highest quality Illumina reads (rq>15) were extracted using trimmomatic (v. 0.36) and mapped versus the reference genomes. Only PE reads where both reads passed the QC step were used for the further steps.

### De novo genome assembly, polishing and annotation

Length-filtered PacBio Sequel reads of NWC_1 were *de novo* assembled with Flye (v. 2.3.1) [45]. We optimised our assembly by setting the minimal read overlap at 3 kb and ran four internal Minimap based polishing rounds (estimated cumulative genome size of 4 Mb). Further, we ran one Arrow polishing step from the SMRTlink (v. 5.0.1.9585) with the Pacbio reads, FreeBayes polishing (v. v1.1.0-56-ga180635; [70]) with the settings “ -F 0.5 --min-coverage 2-p 1” with the Illumina sequences. To test the individual assembly and polishing steps we ran Quast (v4.5) [71]. Further, the NWC_1 genomes were circularized using circlator (v 1.2.1) [72], and all contigs were subjected to three polishing steps using the PacBio reads and Arrow, followed by three additional polishing step using the Illumina reads and FreeBayes.. The filtered ONT reads of NWC_2 were also *de novo* assembled with Flye v.2.3.3 [45] using standard parameters, an estimated cumulative genome size of 8 Mb, and four Minimap polishing iterations. Following the assembly, we manually start-aligned the contigs approximately 200 bp upstream of the *dnaA* gene. Finally, extensive polishing was necessary in the following order: 3x PacBio based arrow polishing, 3x Illumina based FreeBayes polishing, 2x ONT based Racon polishing [73]. To test the individual assembly and polishing steps we ran Quast (v4.5) [71]. Ideel [31] was run to test for an inflated number of pseudogenes, which can serve as an indicator for interrupted ORFs by insertions and deletions. The bacterial genomes and plasmids were annotated with NCBI’s Prokaryotic Genome Annotation Pipeline [74]. All Illumina *de novo* assemblies were done with metaspades and default parameters [32].

### Genome binning

To explore the feasibility of binning, a blobology of the pre-assembled reads from the HGAP assembly was created as previously described [75]. The pre-assembled reads were long and highly accurate (consensus) and taken from HGAP (SmrtLink v. 5.0.1.9585) with the default settings and auto-calculation of the length cutoff. The pre-assembled reads were plotted based on the GC content and coverage as well as the best blast hit (species). The GC content was calculated with EMBOSS infoseq [76], the best alignment and coverage with Minimap2 [77]. Additionally, we calculated the tetranucleotide frequency of the pre-assembled reads [44]. The tetranucleotide frequency and the PC were calculated in R (v3.4.0) using the packages Biostrings and ggplot2.

### Comparative genomics and phylogeny

The GenBank records of completely assembled reference strains of *S. thermophilus* (n=24), *L. delbrueckii* (n=17) and *L. helveticus* (n=34) were downloaded from NCBI RefSeq (as of July 21, 2018). The predicted CDSs of all strains (including our finished MAGs) were used to calculate three maximum likelihood phylogenetic trees using bcgTree [78] (using 100 bootstrap runs while running RAxML [79]). The final output was generated using midpoint rooting in FigTree (v.1.4.3; http://tree.bio.ed.ac.uk/software/figtree/) and modified in Inkscape (v.0.91). The Average Nucleotide Identity was calculated with MUMmer (ANIm) using the jspeciesWS homepage (http://jspecies.ribohost.com/jspeciesws/#analyse, 19.7.2018). To detect variants between two strains, Minimap2 (v.2.10; preset parameters: asm5; [77]) was used to map one assembly to the other. Variants were detected using FreeBayes (v.1.2.0; minimum alternate fraction: 0.1, minimum alternate count: 1). Roary (v.3.12.0) [80] was run using standard parameters to calculate both core and unique genes between two genomes. The CDS of the core and unique genes were compared against the eggNOG 4.5.1 database “bactNOG” (bacteria) and COGs (Clusters of Orthologous Groups) were extracted.

### Taxonomic profiling of NWCs

The species composition of the NWCs was assessed by 16S rRNA amplicon sequencing profiling and analysis of Illumina reads with Metaphlan2 [34]. 16S rRNA amplicon libraries from both NWCs were generated and sequenced on the Illumina MiSeq system using paired-end 250 bp reads at Microsynth (Balgach, Switzerland) according to standard Illumina protocols. PCR amplifications followed a two-step protocol using the Nextera XT DNA library preparation kit. First, 16S rRNA genes were amplified using the standard primers 515F (5’-GTGCCAGCMGCCGCGGTAA) and 806R (5’-GGACTACHVGGGTWTCTAAT) spanning the V4 region [81], followed by the addition of Illumina adapters and indices. The quality of the demultiplexed sequences was inspected using FASTQC (v.0.11.4) and low-quality 3’ ends were trimmed using FASTX Trimmer (v.0.0.14). Subsequent processing steps were performed in Qiime [82]. The trimmed paired-end reads were joined and filtered (Phred quality score of Q20 or higher). Chimeric sequences were removed using USEARCH (v.6.1). OTUs were picked *de novo* and clustered at 99% similarity. The Greengenes database [83] and the BLAST algorithm [84] were used to assign taxonomic identities to the representative sequences of each OTU. Singletons were removed from the OTU table prior to further analyses. In addition to the conventional OTU clustering approach, all joined paired-end sequences were subjected to oligotyping [85]. First, all sequences were trimmed to the same length of 251 bp using Fastx Trimmer. The trimmed reads were subsequently aligned to evaluate the most information-rich nucleotide positions in the alignment using Shannon entropy. To filter out potential sequencing errors, the substantive abundance threshold of each oligotype was set to 100 sequences. The species identification of all oligotypes was verified using BLAST [84]. In addition, the species composition was also assessed using the Illumina raw reads and Metaphlan2 (v.2.7.0; default parameters) [34], and also using the PacBio raw reads and MetaMaps (v.0.1; default parameters) [35].

### Amplification of the slpH locus for *L. helveticus* strain typing

The *L. helveticus* sequence type composition was assessed using a culture-independent strain typing method [36]. Briefly, a 1200-bp region within the *slpH* gene was amplified with the primer pair LHslpF (5’-CAAGGAGGAAAGACCACATGA-3’) and LHslpR (5’-TGTACTTGCCAGTTGCCTTG-3’). The amplicons were fragmented by sonication on a Covaris M220 instrument (Covaris, Brighton, U.K) to obtain 400 bp fragments and subsequently sequenced with the Ion PGM Hi-Q Sequencing kit on an Ion Torrent PGM sequencer (Thermo Fisher Scientific, Baar, Switzerland).

### DNA methylation motif analysis

Prokaryotic methyltransferases methylate the DNA of both bacterial host and plasmids [38]. DNA methylation affects SMRT sequencing by varying the kinetics of the base addition step [86]. To detect any of three major prokaryotic DNA methylation motifs (4-methylcytosine, 5-methylcytosine and 6-methyladenine), a minimum coverage of 250-fold per strand is recommended by PacBio. All DNA methylation motifs were identified using SMRTLink’s Base Modification and Motif Analysis applications (v. 5.0.1.9585). The significance threshold was set to a Benjamini–Hochberg corrected p-value of 0.05 and a quality cutoff of 50.

### Phage identification, annotation and prediction of bacterial host interactions

Similar to a previous study [87], a phage genome database was constructed by downloading all 8056 completely assembled phage genomes from NCBI (as of May 4, 2018). A blastn search of the assembled contigs from NWC_1 and NWC_2 against this database allowed us to identify the most closely related phages, and to subsequently annotate them using the classic RAST pipeline [88, 89]. Prophages were detected and annotated using Phaster [90]. CRISPRFinder [91] was used to identify CRISPR spacers and arrays in all *de novo* assembled NWC genomes, and corresponding spacer sequences were extracted. Next, the assembled phage genomes were specifically searched for matching protospacers with CRISPRTarget [92].

### Statistics and plots

All statistical analyses and plots were performed/created in R (R core team, 3.4.0) using ggplot2 [93]. All circular plots were created with Circos [30].

## List of abbreviations

ANI: Average Nucleotide Identity
CRISPR: Clustered Regularly Interspaced Short Palindromic Repeats
COG: Clusters of Orthologous Groups
EPS: Extracellular Polysaccharides
gDNA: Genomic DNA
HGT: Horizontal Gene Transfer
LAB: Lactic Acid Bacteria
MAGs: Metagenome-assembled genomes
NGS: Next Generation Sequencing
NWC: Natural Whey starter Cultures
ONT: Oxford Nanopore Technologies
PacBio: Pacific Biosciences
PCR: Polymerase Chain Reaction
PE: Paired-End
SNP: Single Nucleotide Polymorphism
WMGS: Whole Metagenome Shotgun

## Declarations

### Ethics approval and consent to participate

Not applicable.

### Consent for publication

Not applicable.

### Availability of data and material

The raw read data is deposited at the NCBI SRA under the Biosample SAMN09703751 and SAMN09580370 for NWC_1 and NWC_2, respectively. The individual genome assemblies are deposited at NCBI Genbank with the following accession numbers:

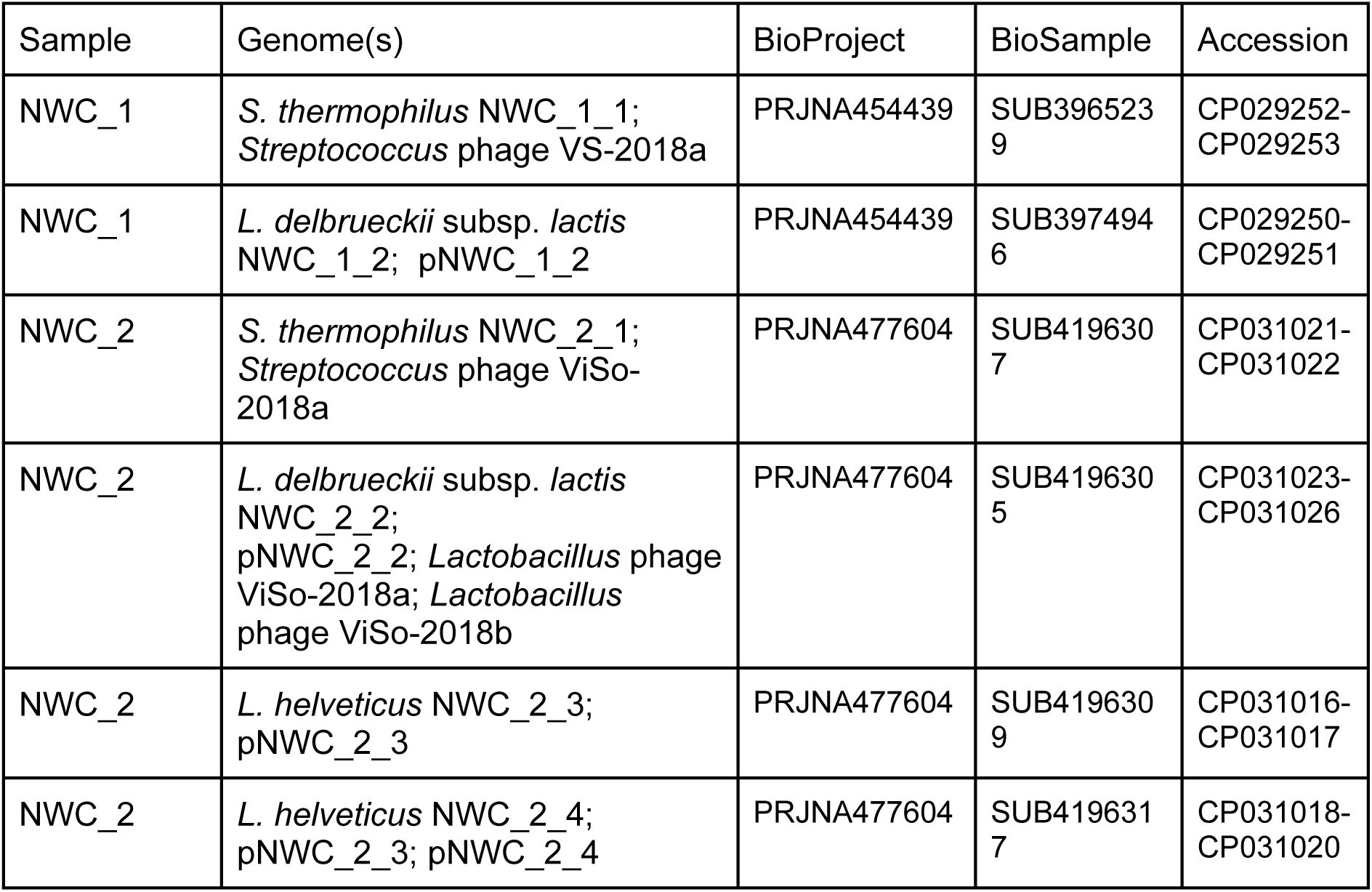

| Sample | Genome(s) | BioProject | BioSample | Accession |
| --- | --- | --- | --- | --- |
| NWC_1 | *S. thermophilus* NWC_1_1;<br>*Streptococcus* phage VS-2018a | PRJNA454439 | SUB396523<br>9 | CP029252-CP029253 |
| NWC_1 | *L. delbrueckii* subsp. *lactis*<br>NWC_1_2; pNWC_1_2 | PRJNA454439 | SUB397494<br>6 | CP029250-CP029251 |
| NWC_2 | *S. thermophilus* NWC_2_1;<br>*Streptococcus* phage ViSo-2018a | PRJNA477604 | SUB419630<br>7 | CP031021-CP031022 |
| NWC_2 | *L. delbrueckii* subsp. *lactis*<br>NWC_2_2;<br>pNWC_2_2; *Lactobacillus* phage<br>ViSo-2018a; *Lactobacillus*<br>phage ViSo-2018b | PRJNA477604 | SUB419630<br>5 | CP031023-CP031026 |
| NWC_2 | *L. helveticus* NWC_2_3;<br>pNWC_2_3 | PRJNA477604 | SUB419630<br>9 | CP031016-CP031017 |
| NWC_2 | *L. helveticus* NWC_2_4;<br>pNWC_2_3; pNWC_2_4 | PRJNA477604 | SUB419631<br>7 | CP031018-CP031020 |

## Acknowledgements

We thank Adithi R. Varadarajan and Mikhail Kolmogorov for constructive input and discussions, and Florian Freimoser for critical feedback on the manuscript.

## Additional Files

**Additional File 1: Table S1.** Overview of raw read statistics for PacBio, ONT and Illumina data. **Table S2:** Assembly statistics for NWC_1 and NWC_2. **Figure S1.** Read distribution of PacBio reads for NWC_1. **Figure S2.** Read distribution of PacBio and ONT reads for NWC_2. **Figure S3.** Phylogenetic tree of completely sequenced *S. thermophilus* strains. **Figure S4.** Phylogenetic tree of completely sequenced *L. delbrueckii* strains. **Figure S5.** Phylogenetic tree of completely sequenced *L. helveticus* strains. **Figure S6.** Binning results for NWC_1. **Figure S7.** Binning results for NWC_2. **Figure S8.** Overview of genome polishing steps. **Figure S9.** Assembly and gene length quality assessment based on Ideel pseudogene plots. **Table S3.** Overview of the total number of transposons identified per bacterial genome. **Table S4.** Results of filtering of reads from 16S rRNA amplicon sequencing. **Table S5.** Analysis of the 16S rRNA V4 amplicon reads using oligotyping. **Figure S10.** Rarefaction curves based on 16S rRNA amplicon sequencing data. **Table S6.** Analysis of the dominant bacterial species in NWC_1 and NWC_2. **Figure S11.** Sequence typing of *L. helveticus* strains in the two NWC samples. **Table S7.** COG categories of *L. helveticus* core and unique genes. **Figure S12.** DNA methylation motif analysis results for NWC_2. **Table S8.** CRISPR arrays identified in NWC_1 and NWC_2. **Figure S13.** Genomic difference of the two *S. thermophilus* strains with respect to copy number of the EPS type VII operon. (PDF)

**Additional File 2:** Overview of the various metrics employed to evaluate each genome polishing step for NWC_2 using QUAST. (XLSX)

