## Supplementary material for "Long read-based *de novo* assembly of low complex metagenome samples results in finished genomes and reveals insights into strain diversity and an active phage system"

### Additional File 1

**Table S1:** Overview of raw read statistics for PacBio, ONT and Illumina data.

**Table S2:** Assembly statistics for NWC_1 and NWC_2.

**Figure S1:** Read distribution of PacBio reads for NWC_1.

**Figure S2:** Read distribution of PacBio and ONT reads for NWC_2.

**Figure S3:** Phylogenetic tree of completely sequenced *S. thermophilus* strains.

**Figure S4:** Phylogenetic tree of completely sequenced *L. delbrueckii* strains.

**Figure S5:** Phylogenetic tree of completely sequenced *L. helveticus* strains.

**Figure S6:** Binning results for NWC_1.

**Figure S7:** Binning results for NWC_2.

**Figure S8:** Overview of genome polishing steps.

**Figure S9:** Assembly and gene length quality assessment based on Ideel pseudogene plots.

**Table S3:** Overview of the total number of transposons identified per bacterial genome.

**Table S4:** Results of filtering of reads from 16S rRNA amplicon sequencing.

**Table S5:** Analysis of the 16S rRNA V4 amplicon reads using oligotyping.

**Figure S10:** Rarefaction curves based on 16S rRNA amplicon sequencing data.

**Table S6:** Analysis of the dominant bacterial species in NWC_1 and NWC_2.

**Figure S11:** Sequence typing of *L. helveticus* strains in the two NWC samples.

**Table S7:** COG categories of *L. helveticus* core and unique genes.

**Figure S12:** DNA methylation motif analysis results for NWC_2.

**Table S8:** CRISPR arrays identified in NWC_1 and NWC_2.

**Figure S13:** Genomic difference of the two *S. thermophilus* strains with respect to copy number of the EPS type VII operon.

**Table S1:** Overview of raw reads statistics for PacBio, ONT and Illumina data.

| Sample | Technology | Raw reads | Raw bases [Gb] | Mean length [bp] | Longest mappable read [bp] |
|---|---|---|---|---|---|
| NWC_1 | PacBio | 379,465 | 1.923 | 5,068 | 19,305 |
| NWC_2 | PacBio | 763,335 | 5.011 | 6,566 | 21,014 |
| NWC_2 | ONT | 407,027 | 1.385 | 3,403 | 118,642 |
| NWC_1 | Illumina PE | 2,132,096 | 1.280 | 300 | 300 |
| NWC_2 | Illumina PE | 1,410,764 | 0.846 | 300 | 300 |

**Table S2:** Assembly statistics for NWC_1 and NWC_2. These include coverage and % of mapping reads of all three sequencing technologies for all assembled genomes. Respective genome features (number of annotated genes, number of 16S rRNA copies) are shown as well.

| Sample | Genome | Size [bp] | Coverage | | | Genes | 16S rRNA | Mapping reads [%] | | |
|---|---|---|---|---|---|---|---|---|---|---|
| | | | Pac Bio | Illu-mina | ONT | | | Illu-mina | Pac Bio | ONT |
| NWC_1 | *L. delbrueckii ssp. lactis* NWC_1_2 | 2,250,954 | 276 | 84 | 0 | 2,286 | 8 | 99.3 | 90.1 | n.a. |
| NWC_1 | pNWC_1_2 | 8,813 | 63 | 72 | 0 | 11 | 0 | | | |
| NWC_1 | *S. thermophilus* NWC_1_1 | 1,899,206 | 560 | 163 | 0 | 2,016 | 6 | | | |
| NWC_1 | *Streptococcus* phage VS-2018a | 39,878 | 365 | 130 | 0 | 55 | 0 | | | |
| NWC_2 | *S. thermophilus* NWC_2_1 | 1,971,439 | 833 | 69 | 160 | 2,108 | 6 | 99.0 | 92.1 | 98.3 |
| NWC_2 | *Streptococcus* phage ViSo-2018a | 15,613 | 7 | 32 | 133 | 15 | 0 | | | |
| NWC_2 | *L. delbrueckii ssp. lactis* NWC_2_2 | 2,269,179 | 273 | 54 | 63 | 2,331 | 8 | | | |
| NWC_2 | pNWC_2_2 | 8,888 | 18 | 89 | 227 | 8 | 0 | | | |
| NWC_2 | *Lactobacillus* phage ViSo-2018b | 41,467 | 43 | 21 | 22 | 86 | 0 | | | |
| NWC_2 | *Lactobacillus* phage ViSo-2018a | 72,362 | 74 | 26 | 155 | 88 | 0 | | | |
| NWC_2 | *L. helveticus* NWC_2_3 | 2,210,811 | 313 | 20 | 95 | 2,385 | 5 | | | |
| NWC_2 | pNWC_2_3 | 22,170 | 1,303 | 163 | 593 | 21 | 0 | | | |
| NWC_2 | *L. helveticus* NWC_2_4 | 2,177,422 | 713 | 34 | 227 | 2,318 | 5 | | | |
| NWC_2 | pNWC_2_4 | 30,538 | 167 | 24 | 133 | 29 | 0 | | | |

**Figure S1:** Read distribution of PacBio reads for NWC_1. a) The cumulative sequencing bases output from the longest to the shortest reads (from right to left) is shown on top. Both x- and y-axes are on a log2 scale. The longest mappable PacBio read was 19,305 bp. b) The read distribution histogram on the bottom was constructed in bins of 50 bp (the x-axis is scaled on a log2 scale).

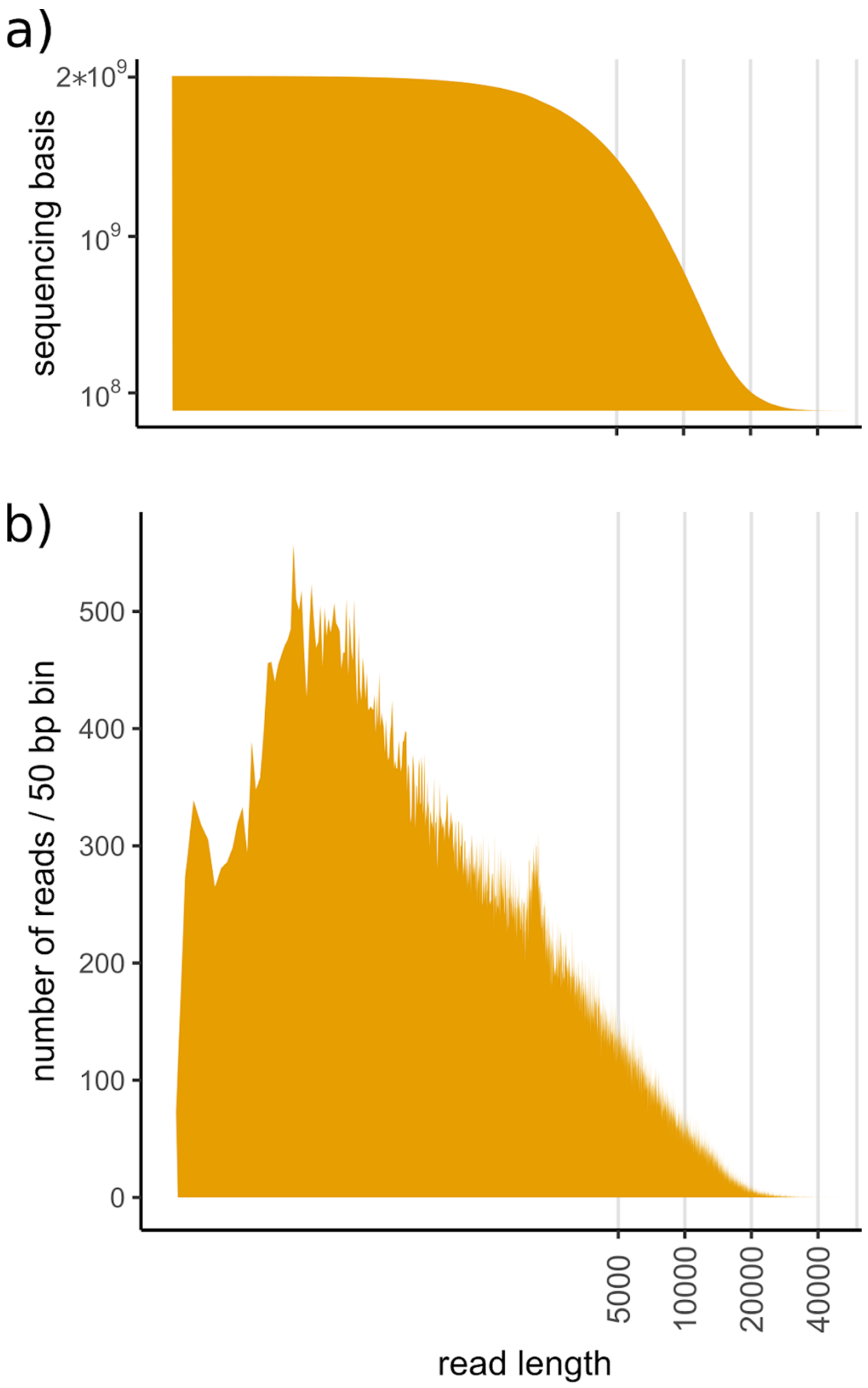

**Figure S2:** Read distribution of PacBio and ONT reads for NWC_2. PacBio data are shown in orange, ONT data in grey. a) The cumulative sequencing bases output from the longest to the shortest reads (from right to left) is shown on top. Both the x and y-axis are on a log2 scale. b) The read distribution histogram on the bottom was constructed in bins of 100 bp (the X-axis is scaled on log2 scale). The longest mappable ONT read is 118,642 bp. The insert visualizes the benefit of adding ONT long reads which are very useful to resolve the dominant genomes. The longest repeats of all completely sequenced *S. thermophilus, L. delbrueckii* and *L. helveticus* strains (NCBI RefSeq) are illustrated as vertical lines.

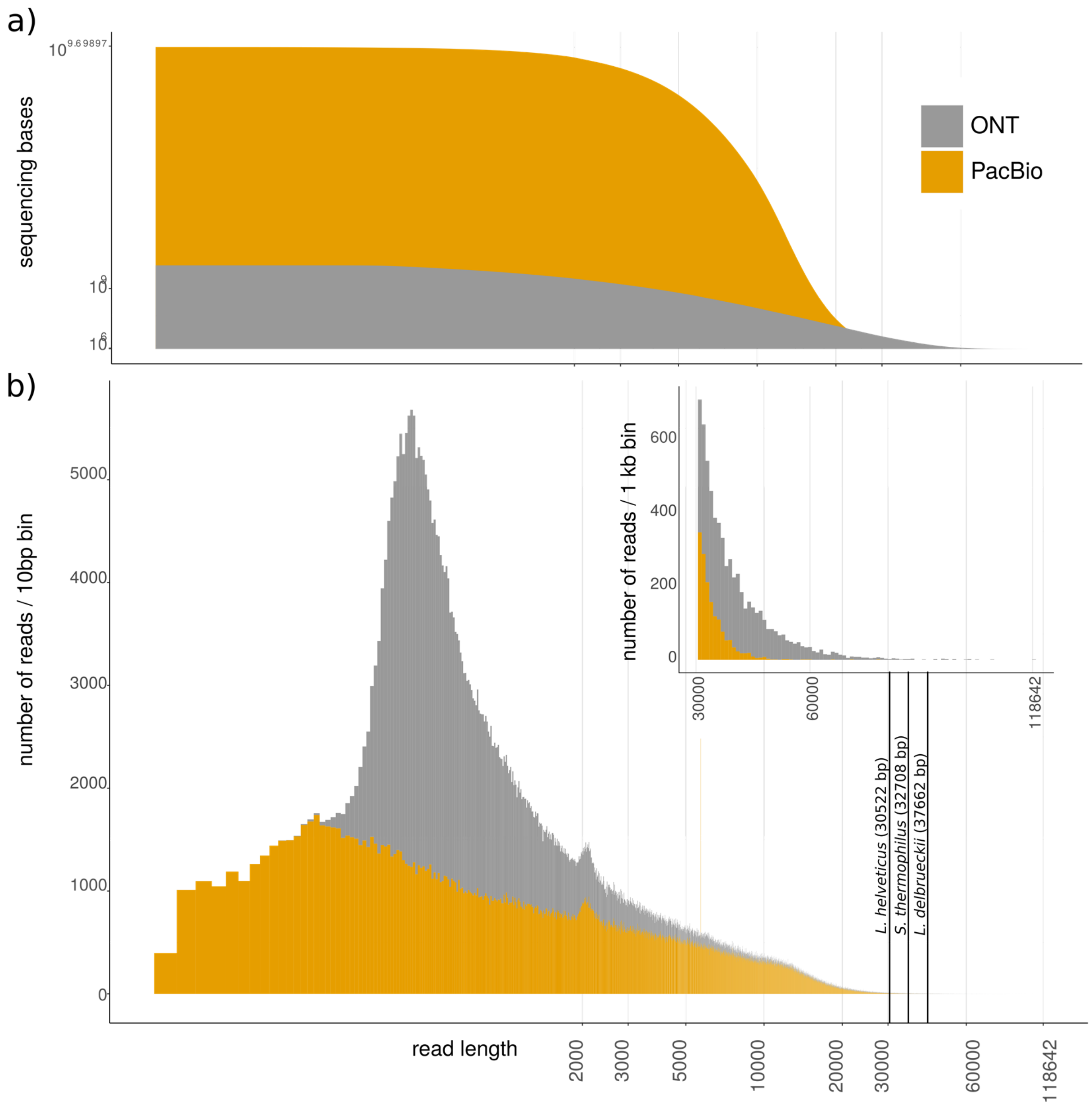

**Figure S3:** Maximum likelihood phylogenetic tree of the two *S. thermophilus* strains NWC_1_2 and NWC_2_2 (in bold) in the context of all currently publicly available and completely assembled *S. thermophilus* strains. The phylogenetic tree was constructed using the core genomes including 107 known housekeeping genes [1]. The bar at the bottom represents the number of amino acid substitutions per site.

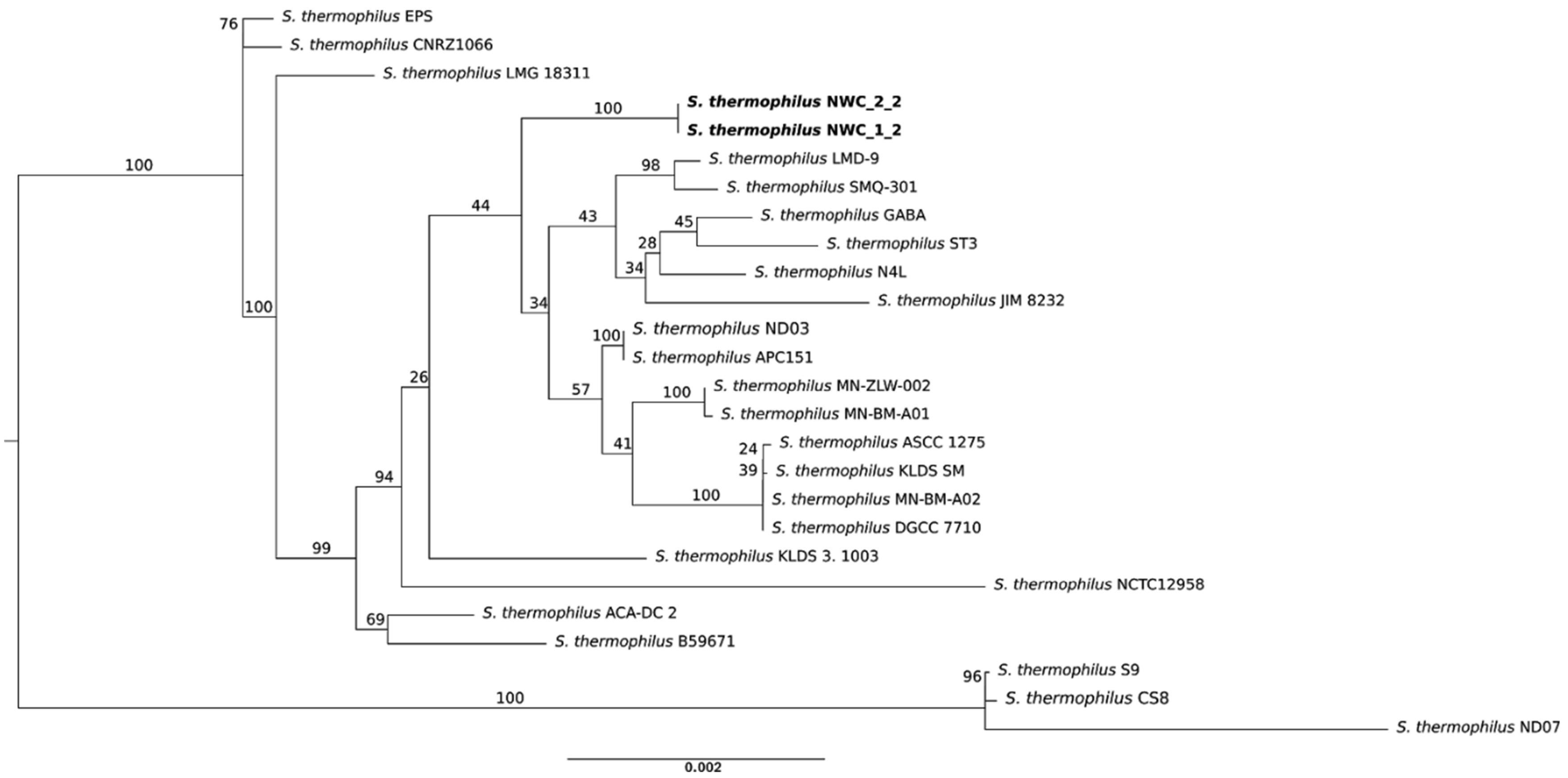


[1] Dupont CL, Rusch DB, Yooseph S, Lombardo M-J, Alexander Richter R, Valas R, et al. Genomic insights to SAR86, an abundant and uncultivated marine bacterial lineage. ISME J. 2011;6:1186–99.

**Figure S4:** Maximum likelihood phylogenetic tree of *L. delbrueckii ssp. lactis* strains NWC_1_2 and NWC_2_2 (in bold) in the context of all currently publicly available and completely assembled *L. delbrueckii* strains. The phylogenetic tree was constructed using the core genomes including 107 known housekeeping genes [1]. The bar at the bottom represents the number of amino acid substitutions per site.

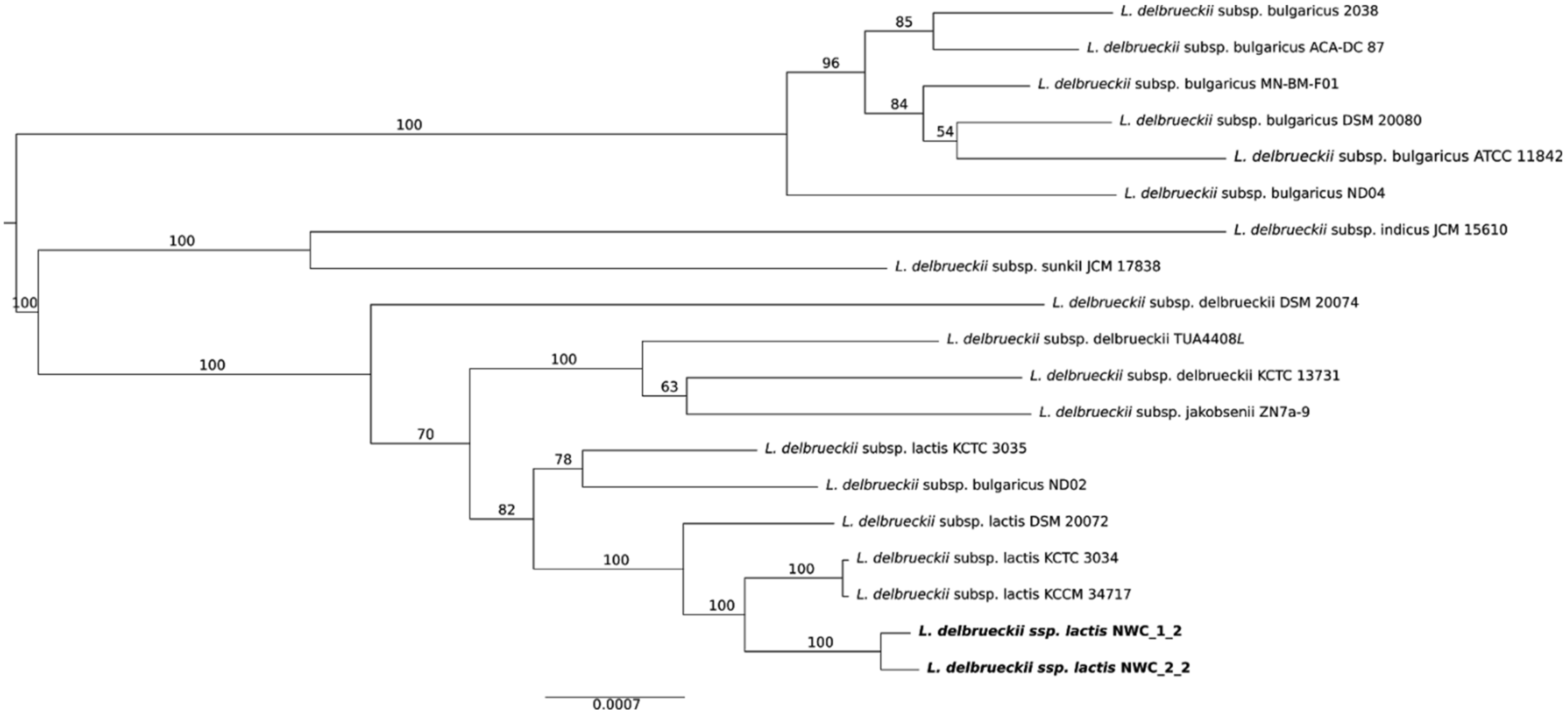


[1] Dupont CL, Rusch DB, Yooseph S, Lombardo M-J, Alexander Richter R, Valas R, et al. Genomic insights to SAR86, an abundant and uncultivated marine bacterial lineage. ISME J. 2011;6:1186–99.

**Figure S5:** Maximum likelihood phylogenetic tree of the two *L. helveticus* strains in NWC_2 (in bold) in the context of all currently publicly available and completely assembled *L. helveticus* strains. The phylogenetic tree was constructed using the core genomes including 107 known housekeeping genes [1]. The bar at the bottom represents the number of amino acid substitutions per site.

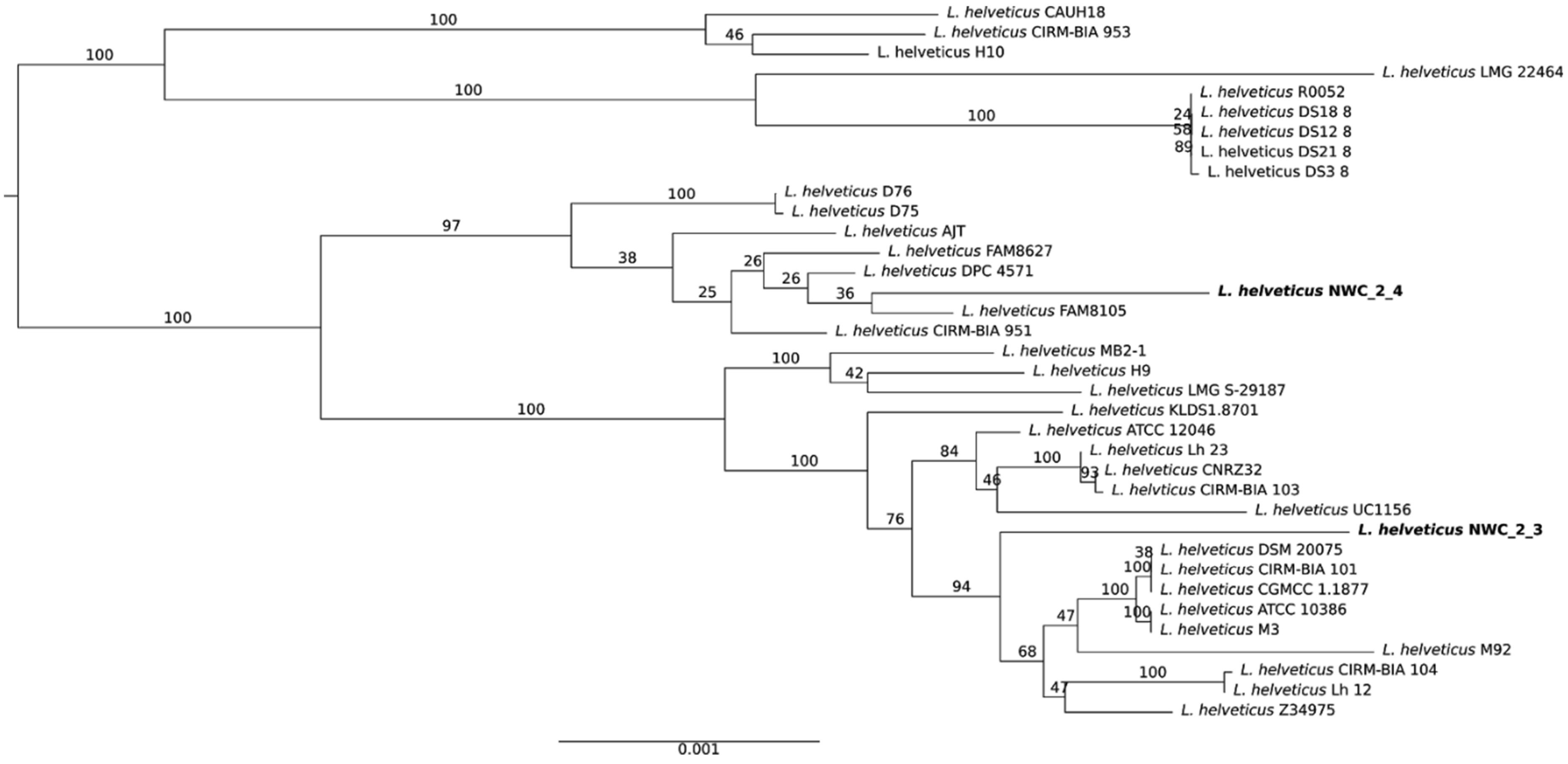


[1] Dupont CL, Rusch DB, Yooseph S, Lombardo M-J, Alexander Richter R, Valas R, et al. Genomic insights to SAR86, an abundant and uncultivated marine bacterial lineage. ISME J. 2011;6:1186–99.

**Figure S6:** "Blobology" (i.e., blobplots or taxon-annotated-GC-coverage plots to visualize the contents of genome assembly data sets) [1] for NWC_1. Binning of the pre-assembled reads (see number of pre-assembled reads per assembled genome in legend) based on GC content, coverage, and PC1 and PC2 of tetranucleotide frequencies, does not allow to completely resolve the different genome compositions in NWC_1. Top left: GC content vs. raw PacBio read coverage. Top right: PC1 vs. PC2 of all tetranucleotide frequencies up to a k-mer size of 5 bp. Bottom left: GC content vs. PC1. Bottom right: GC vs. PC2.

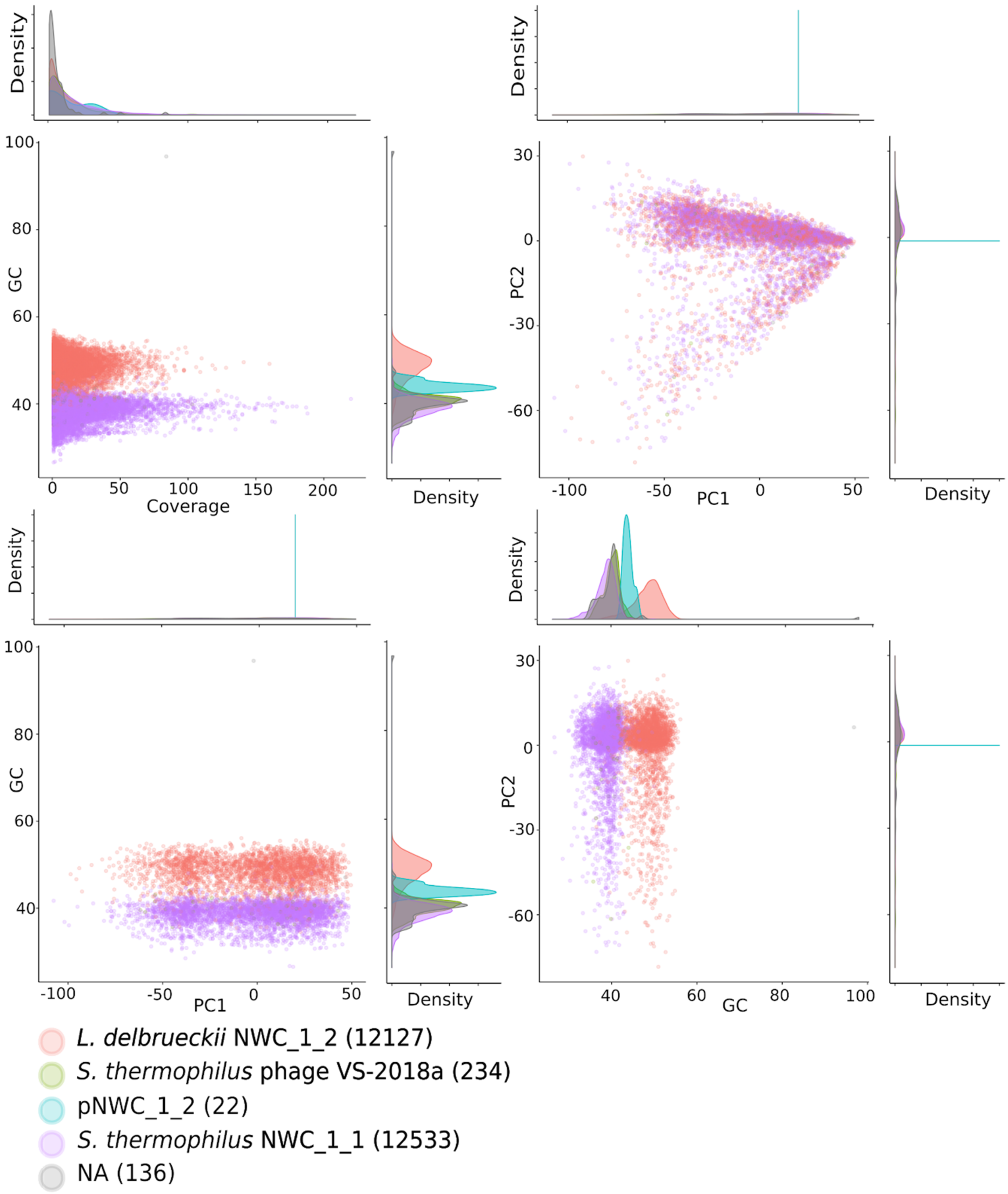


[1] Kumar S, Jones M, Koutsovoulos G, Clarke M, Blaxter M. Blobology: exploring raw genome data for contaminants, symbionts and parasites using taxon-annotated GC-coverage plots. Front Genet. 2013;4:237.

**Figure S7:** "Blobology" (i.e., blobplots or taxon-annotated-GC-coverage plots to visualize the contents of genome assembly data sets) [1] for NWC_2. Binning of the pre-assembled reads (see number of pre-assembled reads per bin in brackets in legend) based on GC content, coverage, and PC1 and PC2 of tetranucleotide frequencies, does not allow to completely resolve the different genome compositions in NWC_2. Top left: GC content vs. raw PacBio read coverage. Top right: PC1 vs. PC2 of all tetranucleotide frequencies up to a k-mer size of 5 bp. Bottom left: GC content vs. PC1. Bottom right: GC content vs. PC2.

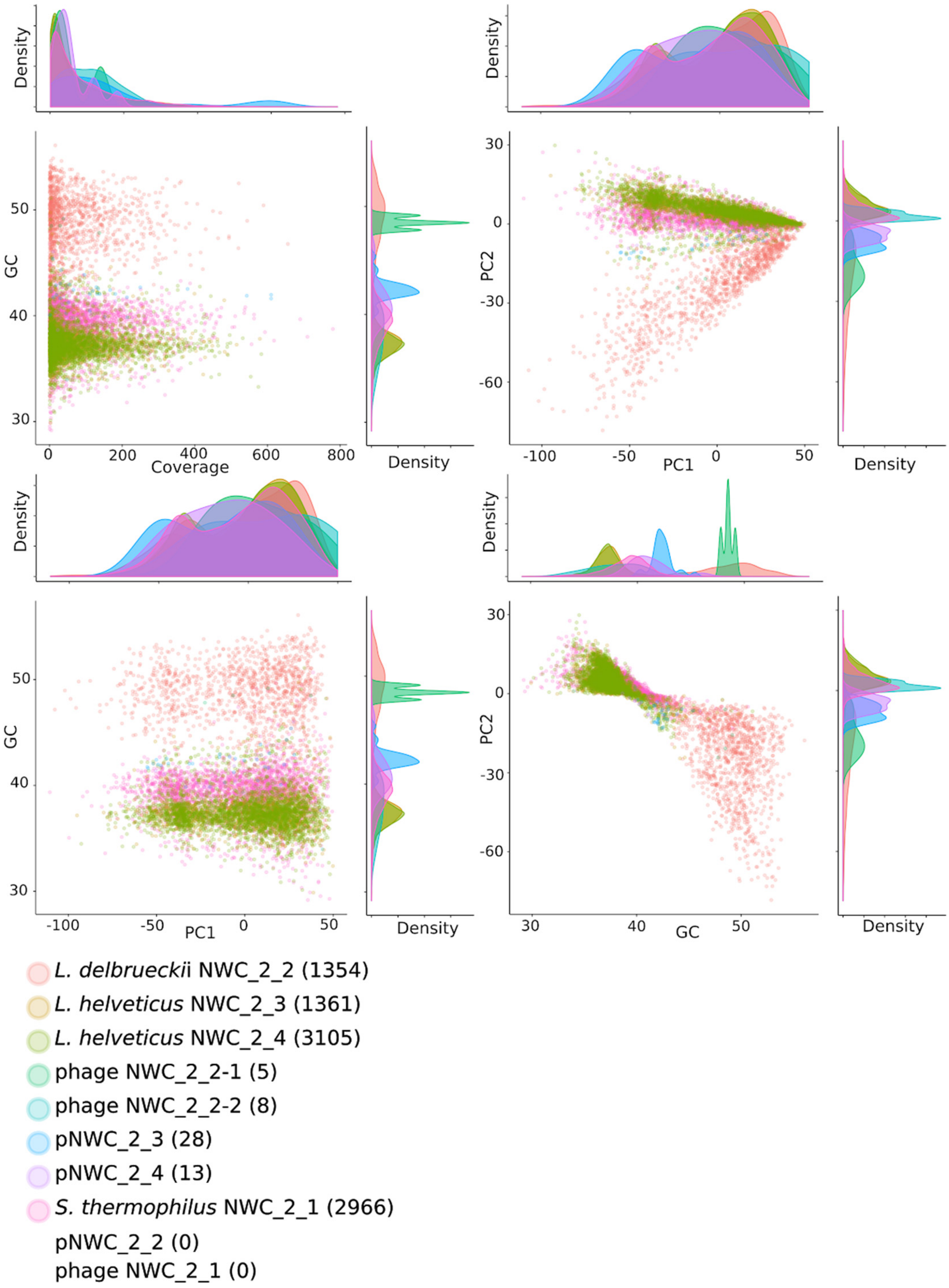


[1] Kumar S, Jones M, Koutsovoulos G, Clarke M, Blaxter M. Blobology: exploring raw genome data for contaminants, symbionts and parasites using taxon-annotated GC-coverage plots. Front Genet. 2013;4:237.

**Figure S8:** Overview of genome polishing steps. Several metrics including number of misassemblies, mismatches per 100kb, pseudogenes and genome fraction indicated that extensive polishing steps after the initial genome assembly were crucial to create finished, high quality genomes. The NWC_1 sample was less diverse and therefore required fewer polishing steps (3x PacBio, 3x Illumina). Extensive polishing of the NWC_2 sample incorporating PacBio, ONT and Illumina data (6x PacBio, 2x ONT, 19x Illumina) resulted in high quality genomes (Fig. 2, Additional File 10). We observed that multiple rounds of polishing of NWC_2 largely improved the quality of the initial genome assembly by removing remaining misassemblies (a), indels, and mismatches (Fig. 2b), and thereby reduced the number of pseudogenes, which can be an indicator for ORFs interrupted by insertions and deletions (c, Additional File 12). For both metagenome samples, we observed that initial rounds of PacBio based polishing with Arrow and Illumina based polishing with FreeBayes [1] removed mainly indels and mismatches (b), yet, only few misassemblies (a). The ONT based polishing with Racon [2] reintroduced a substantial amount of indels, mismatches (b) and misassemblies (a). Nevertheless, the following rounds of FreeBayes and Arrow based polishing indicated that the ONT data most likely resolved previously undetected misassemblies (a), which is illustrated by the steady increase of genome coverage (d). Furthermore, the pseudogene count, which can be taken as a proxy for local genome assembly quality [3], minimized after the final rounds of PacBio and Illumina based polishing (c).

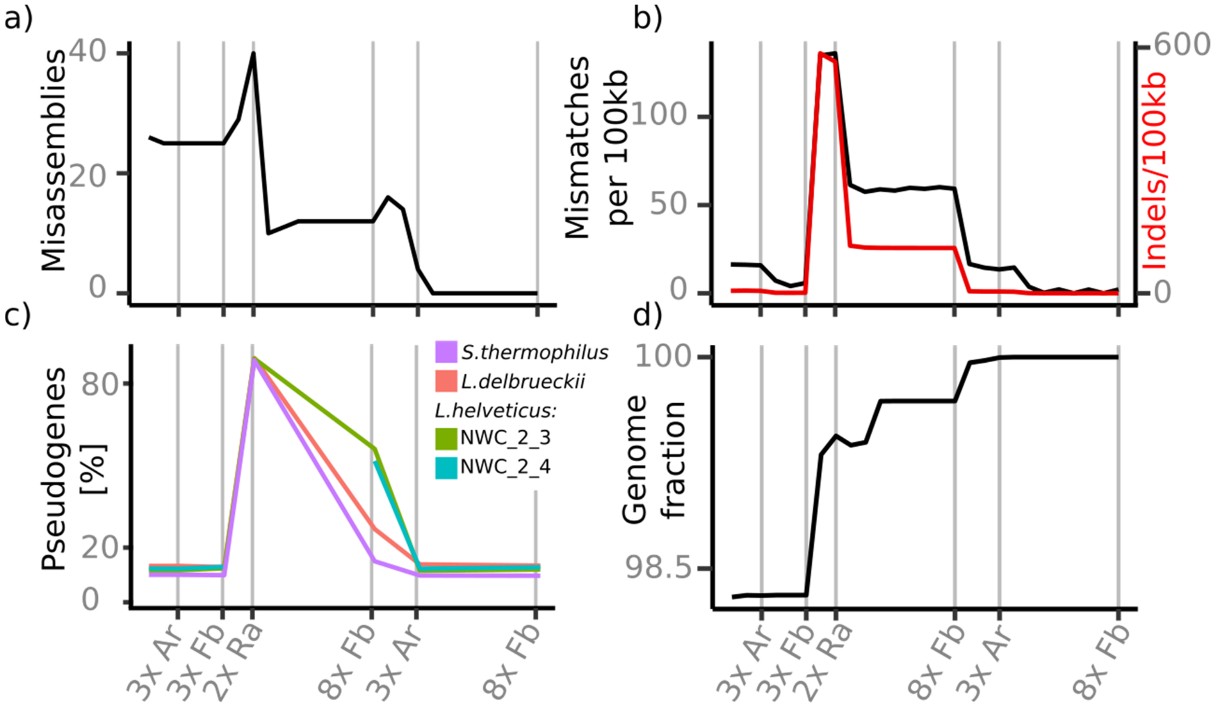


**[1]** Garrison E, Marth G. Haplotype-based variant detection from short-read sequencing. arXiv [q-bio.GN]. 2012. http://arxiv.org/abs/1207.3907. **[2]** Vaser R, Sović I, Nagarajan N, Šikić M. Fast and accurate de novo genome assembly from long uncorrected reads. Genome Res. 2017;27:737–46. **[3]** Watson M. A simple test for uncorrected insertions and deletions (indels) in bacterial genomes. Opiniomics. 2018. http://www.opiniomics.org/a-simple-test-for-uncorrected-insertions-and-deletions-indels-in-bacterial-genomes/. Accessed 12 Jul 2018.

**Figure S9:** Assembly and gene length quality assessment of bacterial genomes assembled from a) NWC_1 and b) NWC_2 based on Ideel pseudogene plots [1]. A large number of short (i.e., interrupted by insertions and deletions) ORFs would appear as a deviation (long tails) from a tight and normal distribution around 1 and indicate an inflated number of pseudogenes. For our assemblies, the deviations from the normal distribution are rather small. Therefore, we conclude that the number of pseudogenes due to sequencing errors is likely low.

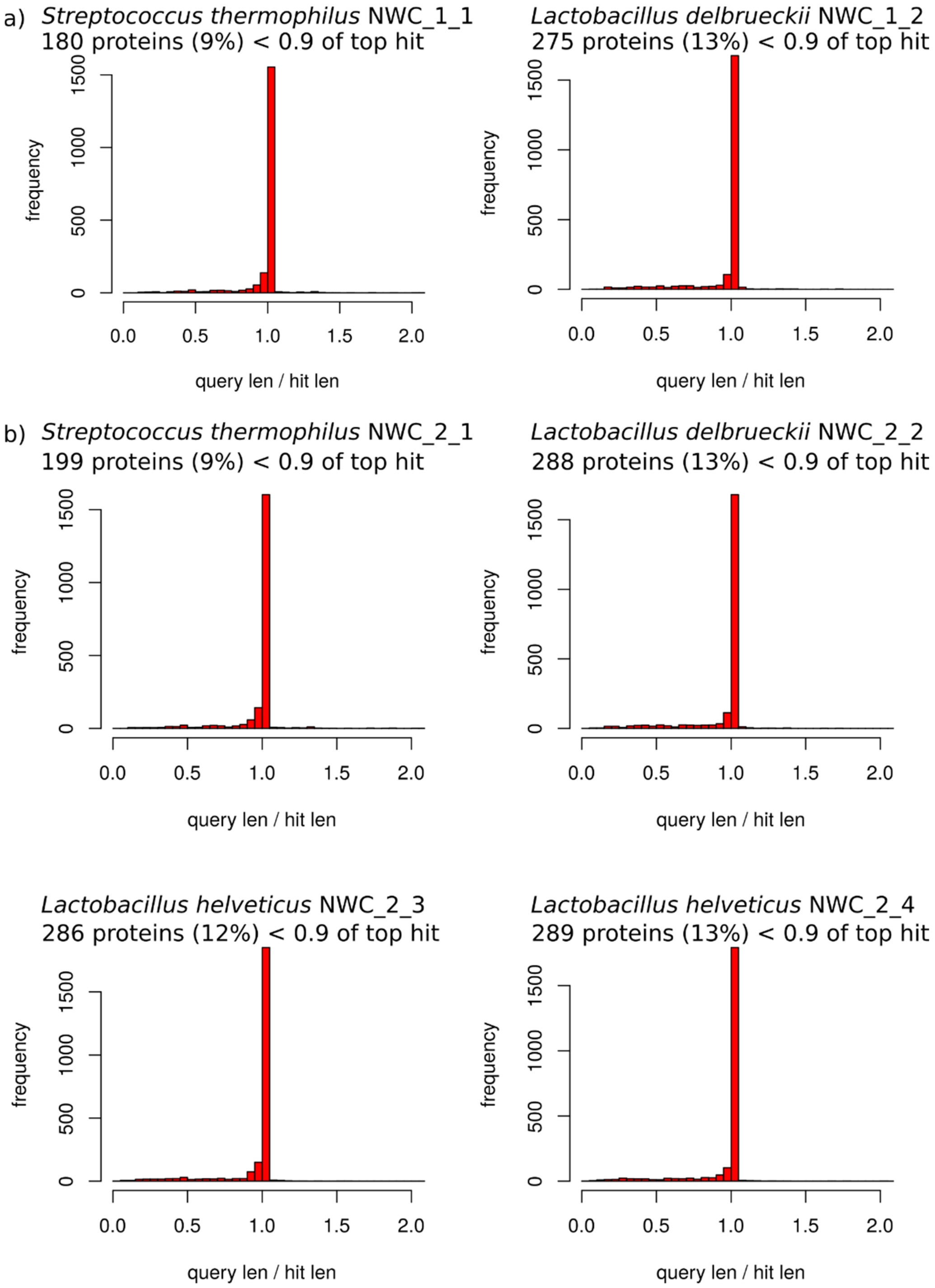


[1] Watson M. A simple test for uncorrected insertions and deletions (indels) in bacterial genomes. Opiniomics. 2018. http://www.opiniomics.org/a-simple-test-for-uncorrected-insertions-and-deletions-indels-in-bacterial-genomes/. Accessed 12 Jul 2018.

**Table S3:** Table with an overview of the overall number of transposases identified in each of the assembled bacterial genomes.

| **Sample** | **Genome** | **No. of transposases** |
|---|---|---|
| NWC_1 | *S. thermophilus* NWC_1_1 | 113 |
| NWC_1 | *L. delbrueckii ssp. lactis* NWC_1_2 | 319 |
| NWC_2 | *S. thermophilus* NWC_2_1 | 121 |
| NWC_2 | *L. delbrueckii ssp. lactis* NWC_2_2 | 291 |
| NWC_2 | *L. helveticus* NWC_2_3 | 467 |
| NWC_2 | *L. helveticus* NWC_2_4 | 406 |

**Table S4:** Filtering of reads from amplicon sequencing (V4 region of 16S rRNA) and further successful assignment to oligotypes [1]. We got a total of 189,583 and 192,631 sequences for NWC_1 and for NWC_2, respectively, of which roughly 94% could be assigned to oligotypes (Additional File 15).

| **Sample** | **No. of raw sequences** | **No. of filtered sequences** | **No. of sequences assigned to oligotypes** | **Percentage (vs raw reads)** |
|---|---|---|---|---|
| NWC_1 | 189,583 | 178,705 | 177,457 | 93.6 |
| NWC_2 | 192,631 | 181,685 | 180,316 | 93.6 |

[1] Eren AM, Maignien L, Sul WJ, Murphy LG, Grim SL, Morrison HG, et al. Oligotyping: Differentiating between closely related microbial taxa using 16S rRNA gene data. Methods Ecol Evol. 2013;4. doi:10.1111/2041-210X.12114.

**Table S5:** Analysis of the 16S rRNA V4 amplicon reads using oligotyping. A conventional OTU clustering approach could only resolve the taxa down to the genus level. However, an oligotyping approach resulted in the delineation of 3 dominant oligotypes overall, which could be identified on the species level, and 6 very low abundant oligotypes, which could be identified either on the species or genus level. The oligotype distribution (relative abundance) in samples NWC_1 and NWC_2 is shown along with their taxon identification and representative sequences. Percentages above 1% are shown in bold. *S. thermophilus* was the dominant species in both samples with a relative abundance of 65.4% in NWC_1 and 45.4% in NWC_2. *L. delbrueckii* was the second most abundant species with a relative abundance of 34.1% in NWC_1 and 24.5% in NWC_2. Two additional oligotypes, which were only present in NWC_2 and only at very low abundances (1.4% and 0.2%), could also be identified as *L. delbrueckii* (>99% similarity). *L. helveticus* made up 0.1% of the community in NWC_1 and 25.6% in NWC_2. Another four oligotypes could only be identified on the genus level as *Lactobacillus spp.; they* made up 0.2% and 2.8% of the total community in NWC_1 and NWC_2, respectively.

| Oligo-type ID | NWC_1 [%] | NWC_2 [%] | Taxon | Sequence |
|---|---|---|---|---|
| GAC | **65.5** | **45.4** | *Streptococcus thermophilus* | TACGTAGGTCCCGAGCGTTGTCCGGATTTATTGGGCGTAAAGCGAGCGCAGGCGGTTTGATAAGTCTGAAGTTAAAGGCTGTGGCTCAACCATAGTTCGCTTTGGAAACTGTCAAACTTGAGTGCAGAAGGGGAGAGTGGAATTCCATGTGTAGCGGTGAAATGCGTAGATATATGGAGGAACACCGGTGGCGAAAGCGGCTCTCTGGTCTGTAACTGACGCTGAGGCTCGAAAGCGTGGGGAGCGAACA |
| ACG | **34.1** | **24.5** | *Lactobacillus delbrueckii* | TACGTAGGTGGCAAGCGTTGTCCGGATTTATTGGGCGTAAAGCGAGCGCAGGCGGAATGATAAGTCTGATGTGAAAGCCCACGGCTCAACCGTGGAACTGCATCGGAAACTGTCATTCTTGAGTGCAGAAGAGGAGAGTGGAACTCCATGTGTAGCGGTGGAATGCGTAGATATATGGAAGAACACCAGTGGCGAAGGCGGCTCTCTGGTCTGCAACTGACGCTGAGGCTCGAAAGCATGGGTAGCGAAC |
| TTA | 0.1 | **25.6** | *Lactobacillus helveticus* | TACGTAGGTGGCAAGCGTTGTCCGGATTTATTGGGCGTAAAGCGAGCGCAGGCGGAAGAATAAGTCTGATGTGAAAGCCCTCGGCTTAACCGAGGAACTGCATCGGAAACTGTTTTTCTTGAGTGCAGAAGAGGAGAGTGGAACTCCATGTGTAGCGGTGGAATGCGTAGATATATGGAAGAACACCAGTGGCGAAGGCGACTCTCTGGTCTGCAACTGACGCTGAGGCTCGAAAGCATGGGTAGCGAAC |
| TTG | 0.0 | **2.4** | *Lactobacillus spp.* | TACGTAGGTGGCAAGCGTTGTCCGGATTTATTGGGCGTAAAGCGAGCGCAGGCGGAAGAATAAGTCTGATGTGAAAGCCCTCGGCTTAACCGAGGAACTGCATCGGAAACTGTTTTTCTTGAGTGCAGAAGAGGAGAGTGGAACTCCATGTGTAGCGGTGGAATGCGTAGATATATGGAAGAACACCAGTGGCGAAGGCGGCTCTCTGGTCTGCAACTGACGCTGAGGCTCGAAAGCATGGGTAGCGAAC |

| | | | | |
|---|---|---|---|---|
| ACA | 0.0 | **1.4** | *Lactobacillus delbrueckii* | TACGTAGGTGGCAAGCGTTGTCCGGATTTATTGGGCGTAAAGCGA<br>GCGCAGGCGGAATGATAAGTCTGATGTGAAAGCCCACGGCTCAAC<br>CGTGGAACTGCATCGGAAACTGTCATTCTTGAGTGCAGAAGAGGA<br>GAGTGGAACTCCATGTGTAGCGGTGGAATGCGTAGATATATGGAA<br>GAACACCAGTGGCGAAGGCGACTCTCTGGTCTGCAACTGACGCTG<br>AGGCTCGAAAGCATGGGTAGCGAAC |
| TCG | 0.0 | 0.3 | *Lactobacillus spp.* | TACGTAGGTGGCAAGCGTTGTCCGGATTTATTGGGCGTAAAGCGA<br>GCGCAGGCGGAAGAATAAGTCTGATGTGAAAGCCCTCGGCTTAAC<br>CGAGGAACTGCATCGGAAACTGTCATTCTTGAGTGCAGAAGAGGA<br>GAGTGGAACTCCATGTGTAGCGGTGGAATGCGTAGATATATGGAA<br>GAACACCAGTGGCGAAGGCGGCTCTCTGGTCTGCAACTGACGCTG<br>AGGCTCGAAAGCATGGGTAGCGAAC |
| ATA | 0.0 | 0.2 | *Lactobacillus delbrueckii* | TACGTAGGTGGCAAGCGTTGTCCGGATTTATTGGGCGTAAAGCGA<br>GCGCAGGCGGAATGATAAGTCTGATGTGAAAGCCCACGGCTCAAC<br>CGTGGAACTGCATCGGAAACTGTTTTTCTTGAGTGCAGAAGAGGA<br>GAGTGGAACTCCATGTGTAGCGGTGGAATGCGTAGATATATGGAA<br>GAACACCAGTGGCGAAGGCGACTCTCTGGTCTGCAACTGACGCTG<br>AGGCTCGAAAGCATGGGTAGCGAAC |
| GCG | 0.1 | 0.1 | *Lactobacillus* spp. | TACGTAGGTCCCGAGCGTTGTCCGGATTTATTGGGCGTAAAGCGA<br>GCGCAGGCGGTTTGATAAGTCTGAAGTTAAAGGCTGTGGCTCAAC<br>CGTGGAACTGCATCGGAAACTGTCATTCTTGAGTGCAGAAGAGGA<br>GAGTGGAACTCCATGTGTAGCGGTGGAATGCGTAGATATATGGAA<br>GAACACCAGTGGCGAAGGCGGCTCTCTGGTCTGCAACTGACGCTG<br>AGGCTCGAAAGCATGGGTAGCGAAC |
| AAC | 0.1 | 0.0 | *Lactobacillus* spp. | TACGTAGGTGGCAAGCGTTGTCCGGATTTATTGGGCGTAAAGCGA<br>GCGCAGGCGGAATGATAAGTCTGATGTGAAAGCCCACGGCTCAAC<br>CATAGTTCGCTTTGGAAACTGTCAAACTTGAGTGCAGAAGGGGAG<br>AGTGGAATTCCATGTGTAGCGGTGAAATGCGTAGATATATGGAGG<br>AACACCGGTGGCGAAAGCGGCTCTCTGGTCTGTAACTGACGCTGA<br>GGCTCGAAAGCGTGGGGAGCGAACA |

**Figure S10:** Rarefaction curve of the 16S rRNA amplicon sequence data. The number of observed OTUs (y-axis) was counted in each sample at different sequencing depths (x-axis). The rarefaction analysis resulted in plateauing curves, which implies that the full diversity was sufficiently sampled.

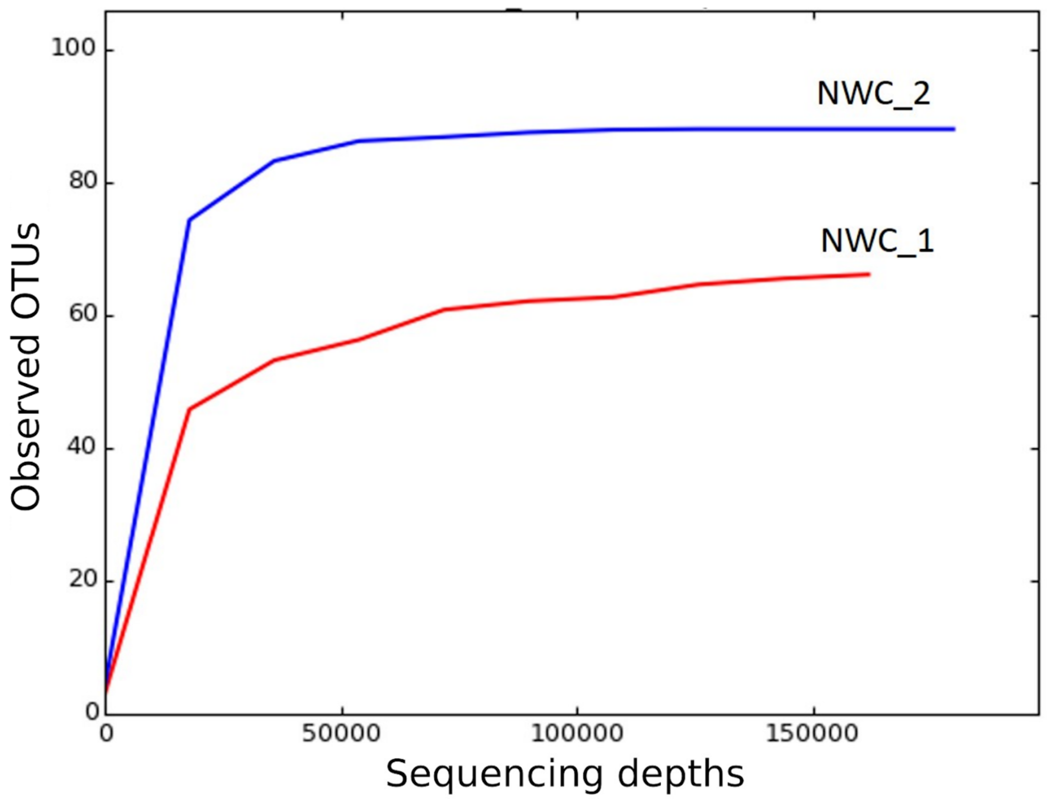

**Table S6:** Analysis of the dominant bacterial species in NWC_1 and NWC_2. We independently assessed the list of dominant bacterial species and strains that we had observed in the two NWCs based on third generation metagenome sequencing data in order to potentially identify additional low abundant taxa. Using the same sample material as input, we thus also carried out a 16S rRNA amplicon-based approach, and explored metagenomic taxon profiling of Illumina and PacBio data using Metaphlan2 [1] and MetaMaps [2], respectively. Numbers represent relative abundances in percentage. MetaMaps can resolve taxon identities down to the strain level and summed relative abundances are given as well as the abundance of the dominant strain(s) in brackets.

| **Sample** | **NWC_1** | | | **NWC_2** | | |
|---|---|---|---|---|---|---|
| **Taxon** | **16S rRNA Oligotyping** | **Illumina Metaphlan2** | **PacBio MetaMaps** | **16S rRNA Oligotyping** | **Illumina Metaphlan2** | **PacBio MetaMaps** |
| ***S. thermophilus*** | 65.4 | 57.6 | 59.3 (ND03: 55.8) | 45.4 | 34.6 | 33.8 (ND03: 31.8) |
| ***L. delbrueckii*** | 34.1 | 42.3 | 32.4 (*subsp. bulgaricus* ND02: 31.0) | 26.1 | 39.2 | 12.2 (*subsp. bulgaricus* ND02: 11.7) |
| ***L. helveticus*** | 0.1 | 0.1 | 0.3 (CNRZ32: 0.3) | 25.6 | 26.2 | 44.5 (DPC 4571: 37.7, CNRZ32: 4.7) |
| **Other** | 0.2 | 0 | 0 | 2.8 | 0 | 0 |
| **Not assigned** | 0.2 | 0 | 8.0 | 0.1 | 0 | 9.5 |

[1] Truong DT, Franzosa EA, Tickle TL, Scholz M, Weingart G, Pasolli E, et al. MetaPhlAn2 for enhanced metagenomic taxonomic profiling. Nat Methods. 2015;12:902–3.

[2] Dilthey A, Jain C, Koren S, Phillippy A. MetaMaps - Strain-level metagenomic assignment and compositional estimation for long reads. 2018. doi:10.1101/372474.

**Figure S11:** Sequence typing of *L. helveticus* strains in the two NWC samples. The different sequence types (STs) were identified using an amplicon-based, culture-independent sequence typing approach [1]. For NWC_1, we detected mainly sequence Type 38 (ST38) with 92% abundance. However, since *L. helveticus* showed an overall abundance of only 0.5 %, it was below the threshold for a complete assembly of dominant genomes using long read metagenomics. For NWC_2, the two dominant sequencing types ST13 (74%) ST38 (19%) corresponded in both abundance (NWC_2_4: 69.9%, NWC_2_3: 30.1%; Fig. 5c) as well as sequence identity to the *slp*H sequences extracted from the assembled *L. helveticus* strains NWC_2_3 and NWC_2_4.

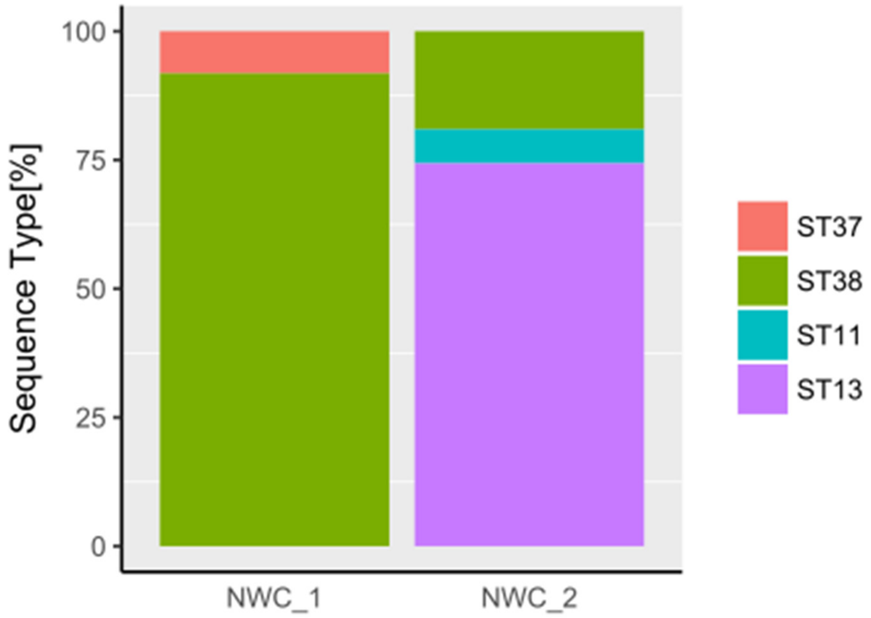


[1] Moser A, Wüthrich D, Bruggmann R, Eugster-Meier E, Meile L, Irmler S. Amplicon Sequencing of the slpH Locus Permits Culture-Independent Strain Typing of *Lactobacillus helveticus* in Dairy Products. Front Microbiol. 2017;8. doi:10.3389/fmicb.2017.01380.

**Table S7:** COG categories of *L. helveticus* core and unique genes. The number of genes associated with COG functional categories for core and unique genes of the two assembled *L. helveticus* strains showed an enrichment for "Nucleotide transport and metabolism (F)" in *L. helveticus* NWC_2_3 and for "Defense mechanisms (V)" in *L. helveticus* NWC_2_4.

| **COG categories** | | **Core genes** | | **Unique genes *L. helveticus* NWC_2_3** | | **Unique genes *L. helveticus* NWC_2_4** | |
|---|---|---|---|---|---|---|---|
| | | Number of genes | % of total genes | Number of genes | % of unique genes | Number of genes | % of unique genes |
| C | Energy production and conversion | 37 | 2.9 | 6 | 1.1 | 6 | 1.1 |
| D | Cell cycle control, cell division, chromosome partitioning | 17 | 1.4 | 2 | 0.4 | 4 | 0.8 |
| E | Amino acid transport and metabolism | 70 | 5.6 | 17 | 3.1 | 21 | 4.0 |
| F | Nucleotide transport and metabolism | 58 | 4.6 | 13 | 2.3 | 1 | 0.2 |
| G | Carbohydrate transport and metabolism | 73 | 5.8 | 9 | 1.6 | 9 | 1.7 |
| H | Coenzyme transport and metabolism | 25 | 2.0 | 4 | 0.7 | 7 | 1.3 |
| I | Lipid transport and metabolism | 33 | 2.6 | 1 | 0.2 | 0 | 0.0 |
| J | Translation, ribosomal structure and biogenesis | 136 | 10.8 | 1 | 0.2 | 4 | 0.8 |
| K | Transcription | 72 | 5.7 | 14 | 2.5 | 12 | 2.3 |
| L | Replication, recombination and repair | 125 | 9.9 | 280 | 50.5 | 259 | 49.3 |
| M | Cell wall/membrane/ envelope biogenesis | 67 | 5.3 | 20 | 3.6 | 13 | 2.5 |

| | | | | | | | |
|---|---|---|---|---|---|---|---|
| N | Cell motility | 0 | 0 | 0 | 0 | 0 | 0 |
| O | Post-translational modification, protein turnover, and chaperones | 42 | 3.3 | 6 | 1.1 | 3 | 0.6 |
| P | Inorganic ion transport and metabolism | 51 | 4.1 | 8 | 1.4 | 10 | 1.9 |
| Q | Secondary metabolites biosynthesis, transport, and catabolism | 2 | 0.2 | 3 | 0.5 | 4 | 0.8 |
| S | Function unknown | 226 | 18.0 | 59 | 10.6 | 55 | 10.5 |
| T | Signal transduction mechanisms | 24 | 1.9 | 2 | 0.4 | 2 | 0.4 |
| U | Intracellular trafficking, secretion, and vesicular transport | 12 | 1.0 | 2 | 0.4 | 1 | 0.2 |
| V | Defense mechanisms | 19 | 1.5 | 2 | 0.4 | 12 | 2.3 |
| Multiple categories | | 11 | 0.9 | 4 | 0.7 | 4 | 0.8 |
| No category | | 158 | 12.6 | 102 | 18.4 | 98 | 18.7 |

**Figure S12:** DNA methylation motifs identified in the *de novo* assembled genomes of NWC_1. The heatmap illustrates the relative abundances (increasing relative abundance from white to black) of the motifs per assembly. We found that the phage shares the same methylation patterns as *S. thermophilus* NWC_1_1. However, we did not observe a DNA methylation motif on the plasmid due to its low coverage, and likely also due to its small size, and thus, fewer potential methylation sites on the plasmid. Therefore, in this case and unlike the plasmids of NWC_2, it could not be matched with its host.

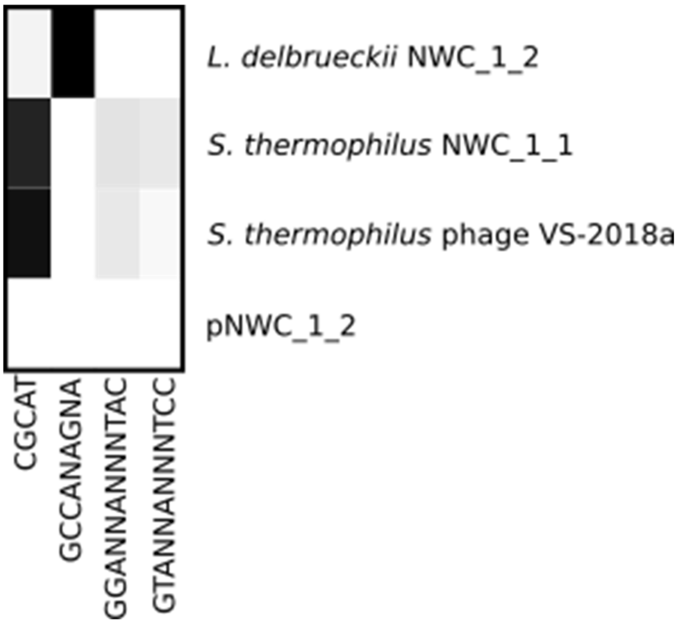

**Table S8:** CRISPR arrays identified in the genomes of strains present in a) NWC_1 and b) NWC_2. The number listed in the respective phage columns indicates the number of matching CRISPR and protospacers (in brackets, we list the number of spacers with more than 3 mismatches).

**a) NWC_1**

| **Genome** | *S. thermophilus* NWC_1_1 | | | *L. delbrueckii ssp. lactis* NWC_1_2 | |
|---|---|---|---|---|---|
| **No. of CRISPR array** | 1 | 2 | 3 | 1 | 2 |
| **Start** | 644,457 | 900,543 | 1,402,794 | 950,430 | 1,163,108 |
| **End** | 646,406 | 901,608 | 1,404,017 | 951,207 | 1,164,559 |
| **length [bp]** | 1,949 | 1,065 | 1,223 | 777 | 1,451 |
| **No. of Spacers** | 29 | 14 | 18 | 11 | 19 |
| **Streptococcus phage VS-2018a** | 3 | 0 | 1 (1) | 0 | 0 |

**b) NWC_2**

| **Genome** | *S. thermophilus* NWC_2_1 | | | | *L. delbrueckii ssp. lactis* NWC_2_2 | *L. helveticus* NWC_2_3 | *L. helveticus* NWC_2_4 |
|---|---|---|---|---|---|---|---|
| **No. of CRISPR array** | 1 | 2 | 3 | 4 | 1 | 1 | 1 |
| **Start** | 328,606 | 1,480,941 | 1,738,880 | 1,739,400 | 1,337,557 | 584,360 | 608,918 |
| **End** | 329,845 | 1,482,872 | 1,739,216 | 1,739,882 | 1,338,603 | 584,655 | 610,428 |
| **Length [bp]** | 1,239 | 1,931 | 336 | 482 | 1,046 | 295 | 1,510 |
| **No. of Spacers** | 18 | 28 | 4 | 6 | 15 | 4 | 22 |
| **Phage1** | 0 | 0 | 0 | 0 | 2 (2) | 0 | 0 |
| **Phage2** | 0 | 0 | 0 | 0 | 0 (1) | 0 | 0 |
| **Phage3** | 1 | 2 (1) | 0 | 0 | 0 | 0 | 0 |
| **Prophage** | 0 | 0 (1) | 0 | 0 | 0 | 0 | 0 |

**Figure S13:** Genomic difference between the *S. thermophilus* strains NWC_1_1 and NWC_2_1. Strain NWC_2_1 harbors two additional copies of an *eps* type VII operon cluster compared to one copy in strain NWC_1_1.

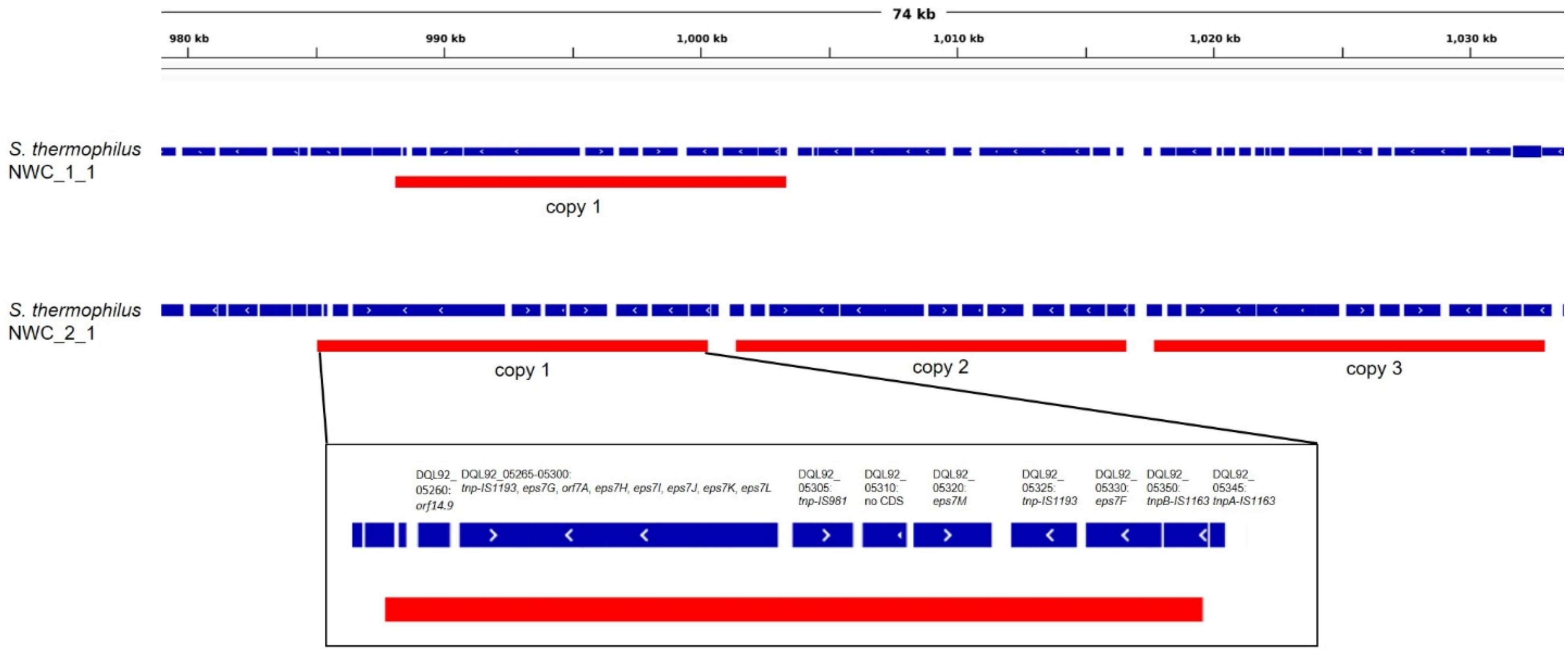
